# A Cross-Species Generative Cell Atlas Across 1.5 Billion Years of Evolution: The TranscriptFormer Single-cell Model

**DOI:** 10.1101/2025.04.25.650731

**Authors:** James D Pearce, Sara E Simmonds, Gita Mahmoudabadi, Lakshmi Krishnan, Giovanni Palla, Ana-Maria Istrate, Alexander Tarashansky, Benjamin Nelson, Omar Valenzuela, Donghui Li, Stephen R Quake, Theofanis Karaletsos

**Author notes:** Corresponding authors: Theofanis Karaletsos and Stephen R. Quake. Authors contributed equally (Sara E Simmonds, Gita Mahmoudabadi, Lakshmi Krishnan).

## Abstract

Single-cell transcriptomics has revolutionized our understanding of cellular diversity, yet our understanding of the transcriptional programs across the tree of life remains limited. Here we present TranscriptFormer, a family of generative foundation models trained on up to 112 million cells spanning 1.53 billion years of evolution across 12 species. By jointly modeling gene identities and expression levels using a novel generative architecture, TranscriptFormer encodes multi-scale biological structure, functioning as a queryable virtual cell atlas. We demonstrate state-of-the-art performance on both in-distribution and out-of-distribution cell type classification, with robust performance even for species separated by over 685 million years of evolution. TranscriptFormer can also perform zero-shot disease state identification in human cells and accurately transfers cell state annotations across species boundaries. As a generative model, TranscriptFormer can be prompted to predict cell type-specific transcription factors and gene-gene interactions that align with independent experimental observations. Developmental trajectories, phylogenetic relationships and cellular hierarchies emerge naturally in TranscriptFormer’s representations without any explicit training on these annotations. This work establishes a powerful framework for quantitative single-cell analysis, and comparative cellular biology, thus demonstrating that universal principles of cellular organization can be learned and predicted across the tree of life.

## Introduction

Diversity in cellular gene expression drives biological complexity at every level, from single-celled organisms responding to their environment and disease, to specialized cells building the tissues and organs of complex organisms, to evolutionary innovations across species. Understanding how these gene expression programs create this cellular variety and how these programs change or persist through evolution is a fundamental challenge in biology.

The explosion of single-cell transcriptomic data, now encompassing hundreds of millions of cells across the tree of life [1–7], presents both opportunities and challenges. While initiatives like CZ CELLxGENE [8] and the Human Cell Atlas [9] have democratized access to harmonized datasets, our ability to compare and interpret data across species boundaries remains challenging. Current approaches for analysis and integration of large single cell corpora [10–13] require using orthologous gene sets to compare across species [14] where distantly related species often share few orthologs, limiting the scope of comparative analysis.

Foundation models have emerged as a powerful paradigm for integrating vast amounts of biological data [15]. Recently, single-cell foundation models, trained on human (scMulan [16], scFoundation [17], scGPT [18]), human and mouse (GeneCompass [19], Nicheformer [20], scPRINT [21]), or multispecies datasets (UCE [22]), have demonstrated that unsupervised learning on transcriptomic data can capture meaningful biological representations, enabling tasks like cell type annotation and gene regulatory network inference. However, these models face key limitations: most are not generative or designed for zero-shot prediction, requiring fine-tuning for each task, and they are typically restricted to human data only. These constraints limit their utility for comparative cellular biology.

Here, we present TranscriptFormer, a family of generative large-scale single-cell foundation models, trained on up to 112 million cells spanning 1.53 billion years of evolutionary history across 12 species (Fig. 1A). TranscriptFormer is a generative autoregressive model capturing the joint probability distribution of genes and their expression levels, enabling it to represent cellular states and facilitate “virtual instruments” for biological inquiry. TranscriptFormer advances the field with a transformer-based architecture coupling gene and transcript heads, expression-aware multi-head self-attention, causal masking, and a count likelihood for transcript-level variability. By training on data from twelve species spanning vertebrates, invertebrates, fungi and protists and incorporating protein sequenced-derived embeddings via Evolutionary Scale Model 2 (ESM-2), which captures protein evolutionary and functional similarity, TranscriptFormer maps genes into a shared, species-agnostic embedding space [15], similar to UCE [22] and scPRINT [21]. This allows TranscriptFormer to learn cell and gene representations that transcend species boundaries while preserving species-specific biology and represents a step forward in unsupervised learning in biology akin to the unsupervised language translation breakthroughs [23] attained by unsupervised pretraining across monolingual language corpora. We assessed how pretraining across a broad corpus of species relates to generalizability in key tasks by evaluating three versions of TranscriptFormer trained on subsets of the corpus: TF-Metazoa (all 12 species), TF-Exemplar (human and 4 model organisms) and TF-Sapiens (human-only data).

**Figure 1.**
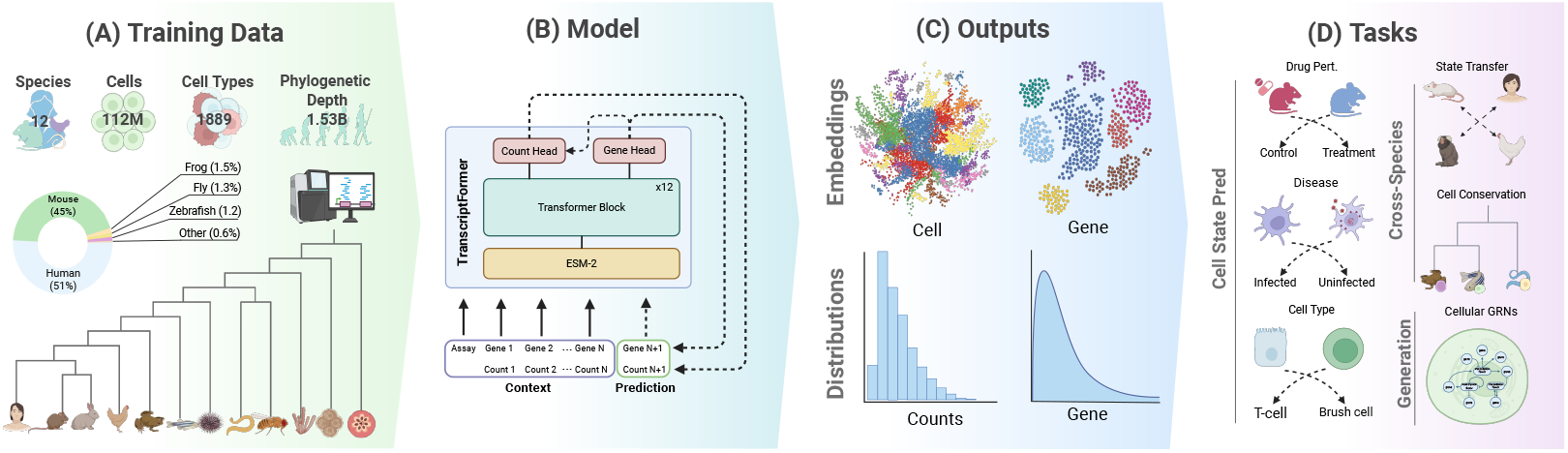
Overview of TranscriptFormer. (A) The largest model TF-Metazoa was trained on 112M cells from single-cell RNA-seq datasets across 12 species spanning 1.53 bn years of evolution. (B) Inputs to the model are cell sentences of protein-coding embeddings of transcripts and their associated raw counts. TranscriptFormer processes inputs to generate representations of expressed genes and cells. (C) Outputs of the model are embeddings and probability distributions at the gene and cell level. (D) Outputs can be used for a variety of downstream tasks, such as cell type classification, disease state prediction, and simulation of gene-gene interactions.

We demonstrate that TranscriptFormer achieves state-of-the-art performance on out-of-distribution (unseen during pretraining) data and zero-shot (without fine-tuning) cell state/type prediction, even in species separated by hundreds of millions of years of evolution. In human perturbation settings, TranscriptFormer accurately detects drug-induced transcriptional states in the Tahoe-100M [24], substantially outperforming benchmark models. We further show that we can reliably transfer drug perturbation state annotations across species without any additional training.

Interestingly, the model reveals emergent biological structure in its cell-level embeddings, such as cell type groupings, developmental trajectories and phylogenetic relationships. Such structures emerge—even in the absence of cell type annotation or metadata in the training data—purely as a consequence of large-scale pretraining on unlabeled transcriptomic data.

Finally, we demonstrate that TranscriptFormer can generate predictions about gene-gene interactions and cell type-specific transcription factors by performing virtual experiments using a novel form of model prompting aligned with the generative AI paradigm. These capabilities suggest that despite the enormous diversity of life, cellular transcriptomes follow learnable, predictable patterns that have been shaped by evolution, principles that can be discovered and understood using AI for biological discovery, with practical utility in tasks related to cell state and cell type prediction.

## Results

### The TranscriptFormer model family

TranscriptFormer is an autoregressive generative model designed to jointly model gene identities and their expression counts as “cell sentences”, capturing the probability of each gene-expression tuple conditioned on preceding tokens (Fig. 1B). Trained via log-likelihood maximization, TranscriptFormer derives biologically meaningful features directly from cellular measurements without reliance on metadata (aside from assay technology identifiers). Similar to UCE [22], gene tokens are represented using ESM-2 [15] protein embeddings (Fig. 1B). Full details are provided in Methods.

The model outputs probabilistic distributions for gene-expression tuples and embeddings for partial or entire cells (Fig. 1C), enabling downstream applications like cell-type classification, disease-state prediction, and cross-species learning (Fig. 1D). Unlike pure embedding-based methods, TranscriptFormer is trained as both a generative and embedding model.

We developed three variants differing by species coverage and evolutionary breadth: **TF-Metazoa**, trained on 112M cells from twelve species (vertebrates, invertebrates, fungus, protist), spanning 1.53 billion years [25] of evolution (Fig. 1A); **TF-Exemplar**, trained on 110M cells from humans and four model organisms (mouse, zebrafish, fruit fly, nematode); and **TF-Sapiens**, trained on 57M human cells. All three models share the same architecture and the same number of parameters in the core transformer layers.

TranscriptFormer was benchmarked against **ESM2-CE** [26], our own baseline model that uses identical input representations but computes fixed-length cell embeddings by averaging ESM-2-derived protein embeddings, omitting the generative modeling component. This approach isolates the specific contribution of TranscriptFormer’s generative architecture.

Evaluations assessed evolutionary generalization, cross-species transfer, and human-specific performance. TranscriptFormer consistently achieved state-of-the-art results, uncovering conserved transcriptional principles across metazoan evolution.

### Cell atlas pretraining enables generalizable cell type and state representations across species

We first evaluated TranscriptFormer’s ability to generalize to species never encountered during training (out-of-distribution species). Using cell atlases from six phylogenetically diverse species, we assessed cell type classification performance, reporting macro F1 scores (Methods). We compared two multispecies TranscriptFormer variants, TF-Exemplar and TF-Metazoa, against the state-of-the-art UCE model [22] and our own baseline ESM2-CE. For calculating F1 scores and uncertainties we apply the linear probing protocol described in Methods. Of these species, human and zebrafish were included in training, while mouse lemur, tropical clawed frog, sea lamprey, and stony coral were completely unseen. Both TranscriptFormer variants maintained robust classification with F1 *>* 0.65 even for stony coral, which diverged approximately 685 million years from humans, while UCE showed significantly lower performance for distant species with F1 ≤ 0.5 (Fig. 2B). Additionally, both variants consistently outperformed ESM2-CE across all species, underscoring the value of training directly on single-cell transcriptomic data rather than protein sequence alone. Overall, TF-Metazoa achieved the highest average F1 score (0.778 ± 0.002), surpassing TF-Exemplar (0.770 ± 0.002) and UCE (0.701 ± 0.002) (Fig. 2C). These results demonstrate that TranscriptFormer effectively captures conserved transcriptional signatures across extensive evolutionary distances.

**Figure 2.**
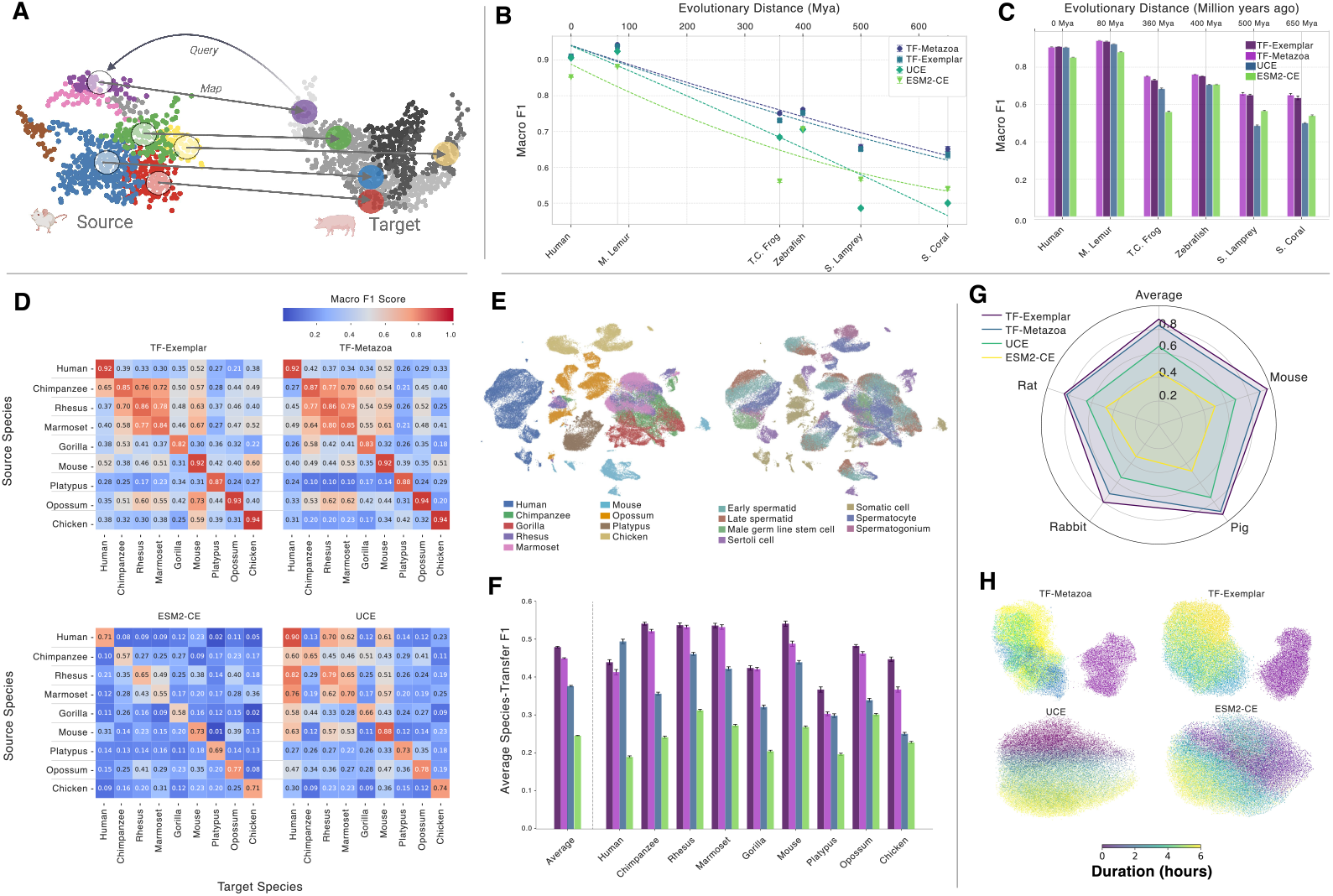
Cross-species transfer learning. (A) Schematic of reference mapping strategy for metadata transfer between species. (B, C) Cell type classification accuracy (macro F1) on cell atlases as a function of evolutionary distance from humans. TF-Metazoa and TF-Exemplar consistently outperform UCE and ESM2-CE, maintaining superior accuracy at larger distances. Species in bold were included in TranscriptFormer pretraining. (D) Transfer matrix of macro F1 scores for testis cell type classification across mammals and chicken, showing performance of each model on all target–reference species pairs. (E) UMAP of TF-Metazoa embeddings from the spermatogenesis dataset, colored by species and cell type, revealing clustering among non-human primates (chimpanzee, gorilla, rhesus, marmoset). (F) Per-species classification accuracy in testis, highlighting TranscriptFormer’s improved generalization to held-out species (e.g., platypus, opossum) compared with UCE and ESM2-CE. Bold species were included in pretraining. (G) Species-averaged transfer F1 scores for LPS perturbation in bone marrow-derived phagocytes from mouse, rat, rabbit, and pig across all models. (H) UMAPs of rat phagocyte embeddings colored by LPS exposure time (0, 2, 4, 6 hours). TranscriptFormer models show clear separation between control (0 hr.) and treated (*>*0 hr.) cells.

We next tested TranscriptFormer’s capability to transfer cell-type annotations between species using the spermatogenesis dataset [27], which spans nine vertebrate species. We evaluated cell-type annotation transfer performance by calculating F1 scores between species pairs. A source species with cell-type annotations is used as a reference to map its annotations to a target species, as illustrated in Fig. 2A and described in more detail in Methods. Generally we found higher F1 scores between closely related species. For example, TF-Metazoa achieved an F1 score of 0.80 for marmoset-to-rhesus transfer, which is comparable to their respective self-transfer scores (0.85 and 0.86) (Fig. 2D). TranscriptFormer models substantially outperformed baselines, with TF-Exemplar achieving the highest species-average F1 score of 0.480 ± 0.002 versus UCE’s 0.377 ± 0.002 (Fig. 2F). Performance gaps increased with evolutionary distance; for chicken-to-mammal transfers (310 million years divergence), TF-Exemplar’s species-average score was 0.448 ± 0.006 compared to UCE’s 0.251 ± 0.004. Notably, TF-Exemplar improved upon ESM2-CE by 91%, emphasizing the substantial benefit gained from incorporating transcriptomic data beyond protein sequences alone. Dimensionality reduction of TF-Metazoa embeddings show extensive overlap among non-human primates (Fig. 2E), reflecting conserved transcriptional programs during spermatogenesis, whereas human cells formed a distinct cluster likely due to technical batch effects (Fig. S2).

Finally, to further assess the cross-species transfer capabilities of TranscriptFormer, we analyzed responses to lipopolysaccharide (LPS) treatment in bone marrow-derived mononuclear phagocytes from mouse, rat, rabbit, and pig [28]. Using the same reference mapping technique for binary classification (control vs. 6-hour LPS treatment), TF-Exemplar and TF-Metazoa achieved F1 scores of 0.925 ± 0.00 and 0.880 ± 0.004, respectively, significantly outperforming UCE (0.740 ± 0.006) and ESM2-CE (0.580 ± 0.007) (Fig. 2G). TranscriptFormer embeddings demonstrated clear separability between control and treatment conditions, in contrast to benchmark models with substantial overlap (Fig. 2H). These results indicate TranscriptFormer robustly captures both stable cellular identities and conserved transcriptional responses to perturbations across diverse species.

### TranscriptFormer infers human cell states across biological contexts

We next assessed TranscriptFormer’s ability to generalize across diverse biological contexts within humans, focusing on cell type classification, disease state prediction, and drug perturbation responses. For cell type classification, we benchmarked TranscriptFormer models (TF-Exemplar, TF-Metazoa, TF-Sapiens) against leading single-cell foundation models: ADIO.Cell [29], scGPT [18], UCE [22], Geneformer [30], scVI [31], and our ESM2-CE baseline. We evaluated the models using the new, previously unpublished component of the Tabula Sapiens 2.0 dataset [32]. We ensured evaluation independence by explicitly holding out the new donors in Tabula Sapiens 2.0 (TSP17-TSP30) from TranscriptFormer pretraining. We calculated tissue-averaged macro F1 scores and uncertainties with the linear probing protocol illustrated in Fig. 3A and described in Methods. TF-Exemplar achieved the highest macro F1 score (0.910 *±* 0.001), closely followed by TF-Metazoa (0.907 *±* 0.001) and TF-Sapiens (0.906 *±* 0.001), which tied UCE (0.906 *±* 0.001) (Fig. 3B). While near-saturation performance occurred on most cell types, TranscriptFormer variants and UCE notably outperformed other models for challenging cell types such as myeloid leukocytes, T-cells, and innate lymphoid cells, maintaining higher classification F1 scores at both the 65% and 95% percentiles (Fig. 3C). Notably, multispecies pretraining did not dilute, and in fact slightly enhanced, performance on human data.

**Figure 3.**
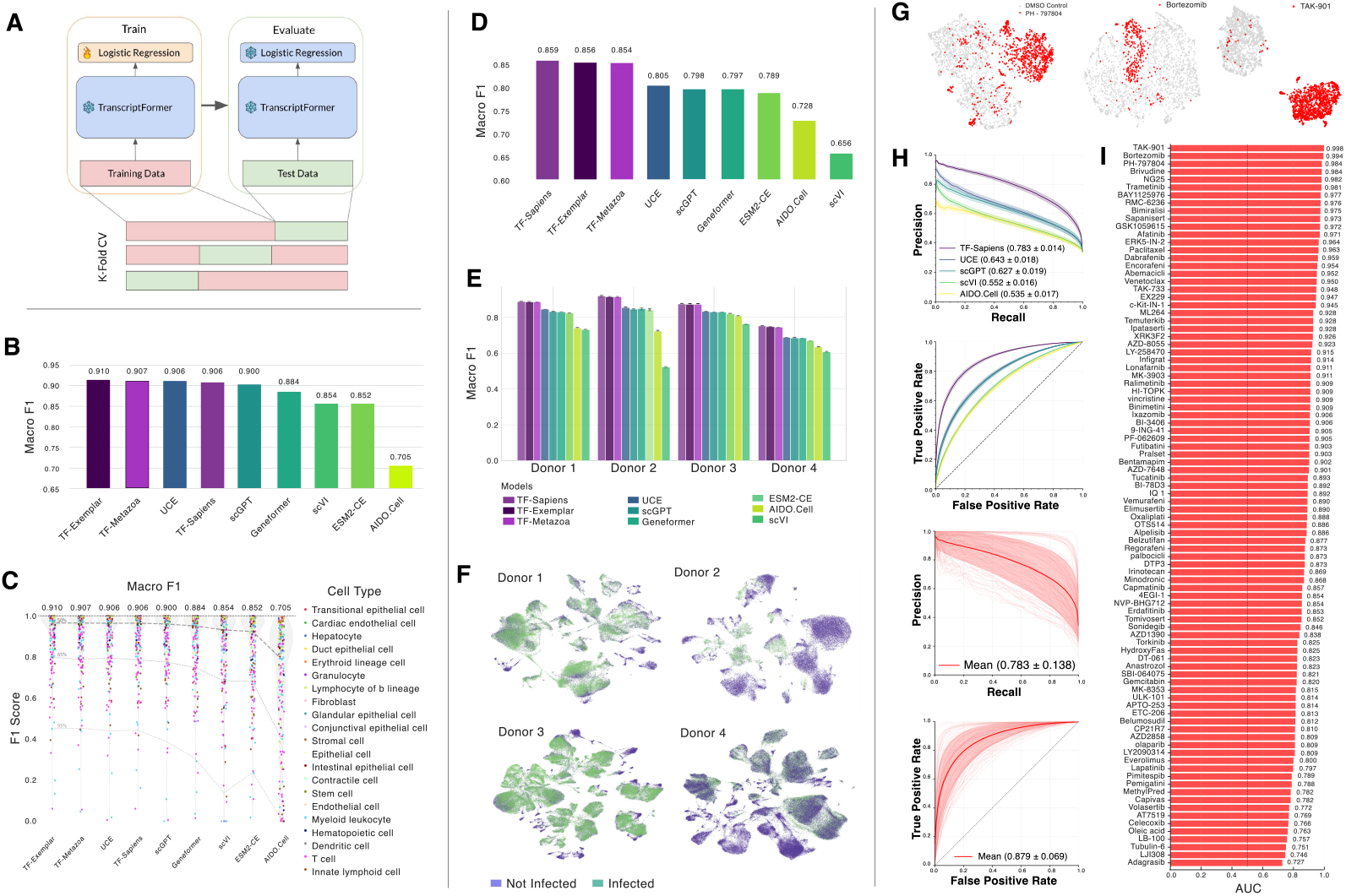
Cell type classification and disease state prediction. (A) Schematic of linear probing protocol used to evaluate TranscriptFormer and benchmark models with k-fold cross-validation. (B) Macro F1 scores for cell type classification on the Tabula Sapiens 2.0 benchmark dataset, averaged across 26 tissues. (C) Per-tissue-per-cell-type F1 scores distributions across models. TranscriptFormer variants and UCE maintain higher F1 scores in the 65th and 95th percentiles compared to other benchmark models. (D) SARS-COV-2 dataset average disease state prediction F1 scores of uninfected and infected cells across models. (E) SARS-CoV-2 dataset disease state prediction F1 scores for each of the four donors across models. (F) TF-Sapiens embedding UMAP of uninfected (purple) and infected (green) cells for each donor. (G) TF-Sapiens embedding UMAP visualizations of control (grey) and treatment (red) on Tahoe-100M drug perturbations. (H) Precision-recall and receiver operator curves (ROC) comparing TF-Sapiens with benchmark models UCE, scGPT, scVI and AIDO.Cell. (I) ROC-AUC binary classification scores for TF-Sapiens across 95 different drug perturbations.

Next, we evaluated disease state classification using a SARS-CoV-2 infection dataset [33] which was not part of the training data, comparing TranscriptFormer variants against baseline models. TF-Sapiens achieved the highest macro F1 score (0.859, uncertainties *<*0.001), with TF-Exemplar (0.856) and TF-Metazoa (0.854) close behind, all significantly outperforming UCE (0.805), scGPT (0.798), and Geneformer (0.797) (Fig. 3D). This trend was consistent across all four donors (Fig. 3E). While cell-type susceptibility could potentially confound disease classification, the comparable cell-type classification performance between TranscriptFormer and baseline models suggests the observed performance differences reflect genuine infection-specific transcriptional changes, as further supported by clear embedding separation between infected and non-infected cells (Fig. 3F).

Finally, we introduced drug perturbations as the last evaluation task, using a sample of the Tahoe-100M dataset [24]. We utilized the same linear probing protocol, framing the task as a series of binary classifications between control (DMSO) and treatment (Methods). TF-Sapiens demonstrated robust detection of drug-induced transcriptional changes across 95 perturbations, achieving a mean AUC of 0.879 *±* 0.007, substantially surpassing UCE (0.779 *±* 0.010), scGPT (0.774 *±* 0.010), and scVI (0.709 *±* 0.008) as shown in Fig. 3H. Analysis of individual drug perturbations revealed considerable variation in classification difficulty, with AUC scores ranging from approximately 0.727 to 0.998 across the 95 compounds tested (Fig. 3I). Several drugs achieved near-perfect classification performance (AUC *>* 0.99), including TAK-901, Bortezomib, and PH-797804, which show clear separable signals in the TF-Sapiens embedding space as visualized by their UMAP projections (Fig. 3G), indicating strong and consistent transcriptional signatures that are readily detectable across cell lines. TranscriptFormer’s superior performance highlights its capacity to detect nuanced biological signals transferable across cellular contexts.

### Emergent hierarchical biological structure in TranscriptFormer cell and gene embeddings

We now explore whether TranscriptFormer learns meaningful biological structures by first examining the models internal representations of genes, within their cellular contexts. TranscriptFormer can compute Contextualized Gene Embeddings (CGEs), which are representations of individual genes, conditioned on all preceding genes in a cell sentence (Fig. 4A, see Methods: CGE).

**Figure 4.**
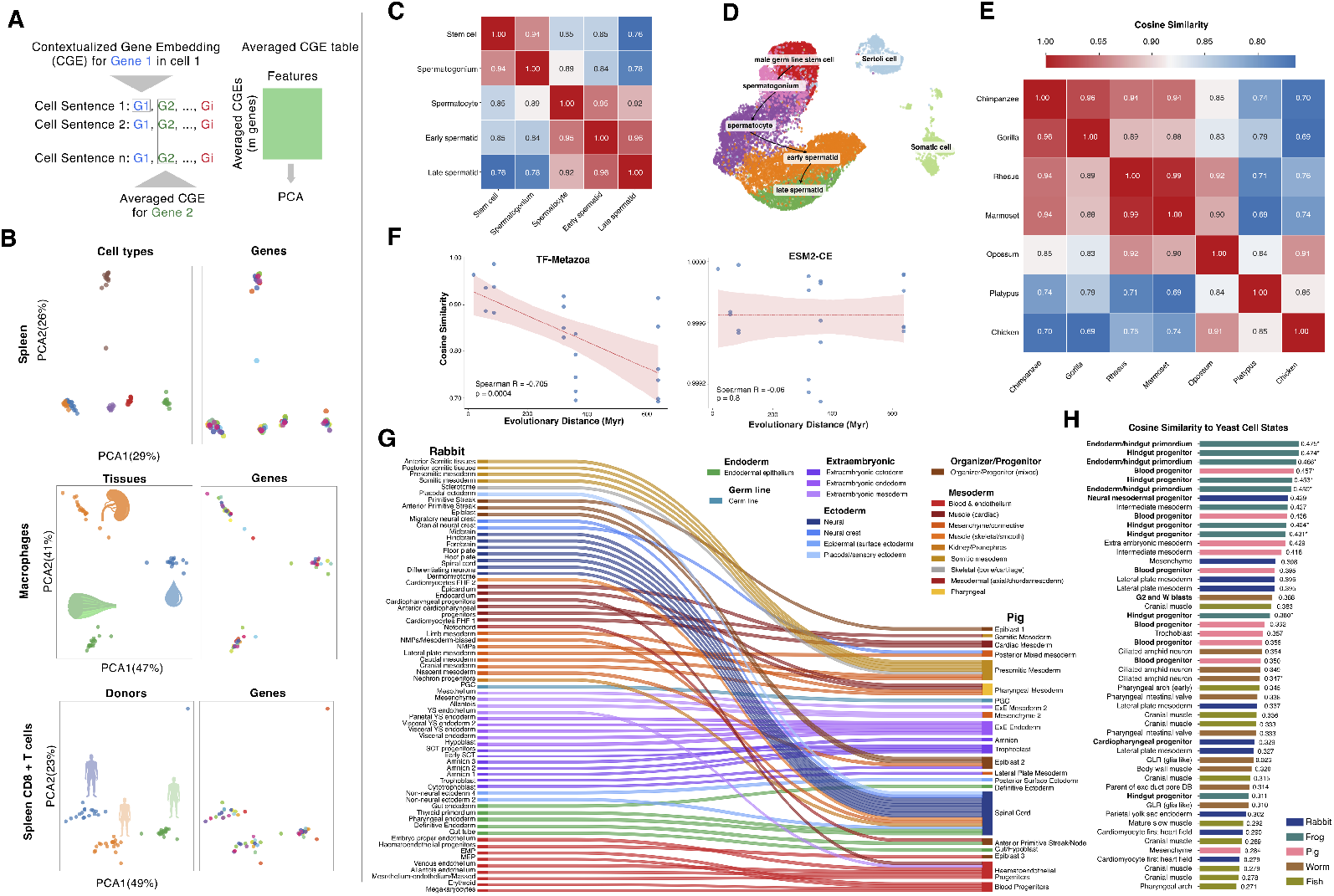
Contextualized Gene Embeddings and cross-species analysis. (A) Schematic representation of the averaged CGEs underlying the PCA plots shown in B. (B) PCA of CGEs from: spleen averaged based on six cell types, macrophages across three different tissues (spleen, muscle and blood), spleen CD8+ T-cells averaged across three different donors (TSP21, TSP25, TSP27). (C) Cosine similarity matrix of germline cell types reveals highest similarity between adjacent differentiation stages (0.89-0.96). (D) UMAP of TF-Metazoa embeddings from human testis cells showing developmental progression from stem cells to mature spermatids. (E) Cosine similarity matrix of pseudo-bulk testis tissue across seven vertebrate species (test set). Red: high similarity; blue: high divergence. (F) TF-Metazoa embedding similarity correlates with evolutionary distance (Spearman’s *r* = -0.705, *p <* 0.001), with closely related species showing greater transcriptional similarity. ESM2-CE embedding similarity shows no correlation with evolutionary distance (Spearman’s *r* = -0.059, *p* = 0.801). (G) Sankey diagram mapping developmental cell atlases across vertebrate species (rabbit-pig), colored by cell class and grouped by germ layer. (H) Cross-kingdom analysis: yeast transcriptional states resemble embryonic progenitor cells across five animal species. Bold indicates progenitor matches and asterisks indicate top match per yeast state.

The CGEs of three tissues taken from Tabula Sapiens 2.0 [32] demonstrate that, despite identical input ESM-2 embeddings for given genes, they cluster primarily by cell type. This clustering pattern persists across all three tissues, validating the model’s ability to capture cell type-specific context across a diverse range of genes (Fig. 4B (top), Fig. S3), without having access to cell type labels. Further analysis of shared cell types between tissues reveals that CGEs incorporate the subtle variations in transcriptomic profiles of identical cell types across different tissues in a zero-shot manner, highlighting the context-sensitivity of the model’s gene representations (Fig. 4B (middle), Fig. S4).

TranscriptFormer also exhibits sensitivity to donor-specific transcriptomic variations, with CGEs from the same cell type and tissue separating based on donor (Fig. 4B (bottom), Fig. S5). We further show that donor effects are stronger than batch effects, however we cannot completely rule out batch effects (Fig. S7). This finding underscores TranscriptFormer’s capacity to detect inter-individual variability perhaps in part stemming from biological factors such as genetic polymorphisms or environmental exposures. Moreover, we used variance partitioning analysis to quantify the contribution of cell type, tissue and donor contexts to CGEs. We show that cell type information dominates the CGE space (explaining *>*95% of variance in PC1 and PC2, after accounting for tissue and donor), while tissue and donor information contribute as secondary factors (explaining *<*2% in PC1 and *<*7% in PC2, after accounting for the other two variables) with this hierarchy remaining consistent across various gene selection criteria (see Methods, Fig. S6). These results demonstrate TranscriptFormer’s ability to encode cell type, tissue type, and donor information in a unsupervised manner.

To assess whether TranscriptFormer cell-level embeddings capture higher-order biological processes, we examined their structure, finding evidence of emergent organization across multiple dimensions. We found that cell embeddings from the spermatogenisis dataset [27] capture known cell types and male germ line differentiation relationships, with adjacent cell types having the highest cosine similarity in embedding space, reproducing the canonical order of spermatogenic stages (Fig. 4C-E).

We also observed that TF-Metazoa cell embeddings encode phylogenetic structure across species as pairwise cosine similarity decreases with evolutionary distance (Spearman’s *r* = −0.705; *p* = 0.004) (Fig. 4F (left)). This pattern emerges despite the fact that TranscriptFormer was not trained on any of the species, showing that the model has strong generalization capabilities. In contrast, embeddings from ESM2-CE show no significant correlation of cosine similarity with evolutionary distance (Spearman’s *r* = −0.059; *p* = 0.801) (Fig. 4F (right)).

Next, we curated a 1.6M cell multispecies developmental cell atlas to find the most similar cells across species (Methods). Using TranscriptFormer embeddings, we recovered known conserved cell types across species of varying evolutionary distance. For example, mapping between closely related mammals, pig and rabbit, revealed correspondence of broad cell classes involved in key developmental systems: primordial germ cells (PCG), extraembryonic tissues (extraembryonic endoderm, mesoderm and ectoderm), blood & endothelium (blood progenitors, haematoendothelial progenitors), presomitic mesoderm, endodermal epithelium and neural cells (Fig. 4G).

Comparing broad cell classes across more distantly related vertebrates, clawed frog and zebrafish, showed cell classes mapping in mesodermal (skeletal muscle, blood & endothelium) and ectodermal (neural, neural (retina/CNS)) cells (Fig. S8). Zebrafish and rabbit (429 MYA) cell mapping revealed cell types involved in some vertebrate developmental programs matched including mesodermal (blood & endothelium) and ectoderm (differentiating neurons, hindbrain, floor plate, migratory neural crest) (Fig. S9). For example, rabbit migratory neural crest mapped to zebrafish cranial neural crest, neural crest + pigment cell progenitor, and iridophores (iridescent pigment cells), all known neural crest derivatives in vertebrates [34]. The zebrafish swim bladder cells mapped to cells in the frog gut primordium and rabbit gut tube. The swim bladder is known to derive from the gut primordium in vertebrates [35]. Hematopoiesis, the development of blood cells, showed consistent mapping across all vertebrate species analyzed, confirming its deep evolutionary conservation in vertebrates [36]. This cross-species cell type mapping demonstrates that TranscriptFormer embeddings capture some known developmental biology programs.

The deepest evolutionary comparison between *C. elegans* and clawed frog revealed some surprising mappings despite representing one of the major splits in animal development (protostome vs deuterostome, ∼600 MYA). Basic muscle and neural cell types consistently matched (Fig. S10), suggesting these fundamental cell types emerged before these major animal groups diverged, consistent with known evolutionary history of neurons and muscle cell families [14]. Interestingly, we found that coelomocytes, macrophage-like cells, in *C. elegans* mapped to migrating myeloid progenitor cells in frog that give rise to macrophages, lending support to characterization of coelomocytes as macrophage-like immune cells [37]. We observed mapping between secretory glands: cells from the frog cement gland primordium mapped to *C. elegans* pharyngeal gland. Both structures secrete mucin proteins, demonstrating that despite vast evolutionary distances, specialized secretory cells might share similar gene expression patterns and could be an example of convergent evolution.

Finally, we attempted to map cell states across kingdoms from fungi to whole-body developmental cell atlases of 5 animals, spanning more than one billion years of evolution. We found that yeast cell states grown under ten different environmental conditions were most similar (9/10 top hits) to embryonic progenitor cell states in clawed frog (endoderm/hindgut primordium, hindgut progenitor, intermediate mesoderm progenitor), pig (blood progenitor), and rabbit (neural mesodermal progenitor) (Fig. 4H). This suggests that the transcriptional states of unicellular organisms like yeast resemble the flexible, undifferentiated states of animal multipotent cell states rather than fully differentiated cell types.

A fundamental question in machine learning for biology is whether models can discover biologically meaningful structure from unsupervised training. Despite training TranscriptFormer without any cell type annotations, developmental stage labels or phylogenetic information, the model learned hierarchical biological organization from molecular to evolutionary scales.

### Using TranscriptFormer as a virtual instrument via prompting

TranscriptFormer models the joint distribution of genes and expression levels across cells, similar to a generative language model. We exploited this generative capability of the model to utilize it as a virtual instrument, and demonstrated how to prompt the model to represent complex inquiries into cellular biology that would otherwise be tackled by compiling datasets laboriously.

Our first prompt asked the model to predict interactions between transcription factors and other protein coding genes using its generative interface. We obtained point-wise conditional mutual information (PMI) estimates between these transcription factors-gene pairs (see Methods), which we used to identify high-confidence interactions (adjusted *p*-value ≤ 0.05) and cross-reference with known biological relationships documented in the STRING database v12.0 [38]. PMI was chosen because it highlights specific co-occurrences between genes and transcription factors while correcting for genes with high baseline expression.

TF-Sapiens accurately recovered key regulatory targets for six transcription factors with diverse biological functions (Fig. 5A). For cell cycle regulation, TranscriptFormer identified canonical targets for three key regulators: E2F8 (87/227 predictions validated in STRING [38]), FOXM1 (105/224 validated), PTTG1 (70/106 validated) and MYBL2 (54/87 validated), correctly linking them to replication factors (CDT1), mitotic regulators (TPX2, HJURP, KIF18B; [39, 40] [41, 42], [43]), and spindle checkpoint genes (BIRC5, PBK) respectively (Fig. 5A). MYBL2 is correctly linked to late cell-cycle regulators (CDC45, BIRC5), reflecting its cooperation with FOXM1 in mitotic transcription [44]. The model similarly captured lineage-specific regulations, identifying B-cell markers CD19 and CXCR5 as SPIB targets (9/23 validated) supporting its function in immune lineage commitment [45, 46]. Finally, UHRF1 (54/90 validated) is associated with replication stress and checkpoint factors like CHEK1 and TIMELESS, consistent with its epigenetic role in coordinating DNA methylation and genome stability [47, 48].

**Figure 5.**
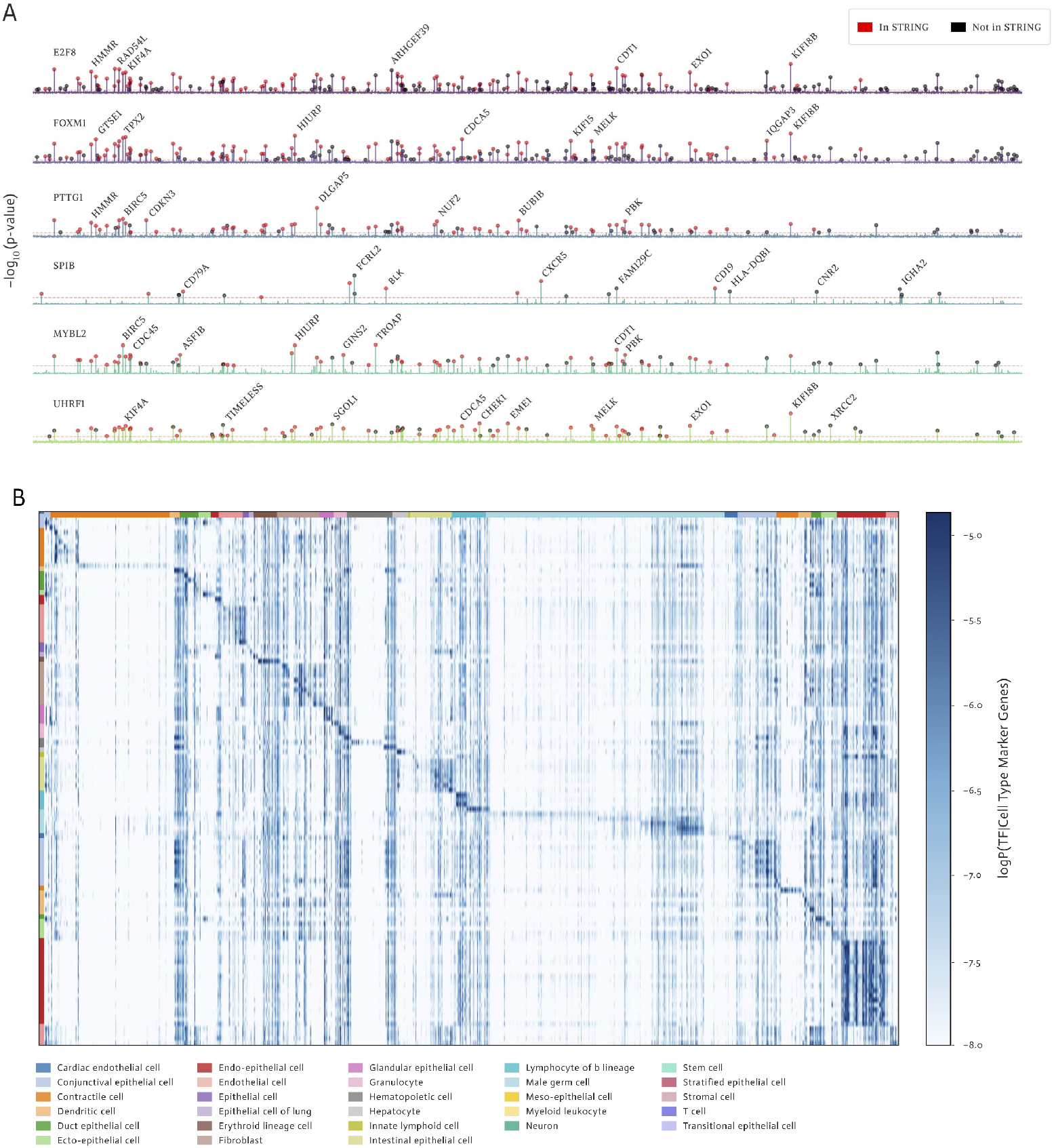
Cell type classification and disease state prediction. Predicting genegene relationships and cell type-specific transcription factors. (A) Point-wise conditional mutual information (PMI) spectra for selected transcription factors, computed using Tran-scriptFormer’s autoregressive architecture showing gene-level associations, with genes annotated in STRING v12.0 shown in red. (B) TranscriptFormer-predicted gene expression log-probabilities for transcription factors across 112 human cell types from Tabula Sapiens 2.0. TF-Sapiens generates known structural features: a diagonal pattern of cell type-specific transcription factors and vertical bands of broadly expressed regulators.

Next we created more complex prompts by conditioning the model on a set of marker genes corresponding to specific human cell types from the Tabula Sapiens 2.0 [32] dataset. We generated cell-type conditioned transcription factor-gene interaction probability distributions for all 112 known human transcription factors. Our model-generated heatmap (Fig. 5B) recapitulates structural features observed in empirical data from Tabula Sapiens 2.0. We employed a methodology that parallels the approach used in the Tabula Sapiens 2.0 study, with the key distinction that transcription factor-gene interactions were generated by the model rather than measured experimentally from gene expression. We arranged transcription factors according to the cell type for which they showed the highest probability of expression, creating a visualization that mirrors Figure 3A in the Tabula Sapiens 2.0 publication. Our generated heatmap exhibits remarkably similar global structures: vertical bands corresponding to ubiquitously expressed transcription factors that function across diverse cell types, and a distinctive diagonal trace representing cell type-specific transcription factors with highly specialized expression patterns. While different cell type annotations between our evaluation data and the original Tabula Sapiens 2.0 dataset prevent a one-to-one comparison, we did recover several key transcription factors and broad cell types associations mentioned in [32], including IKZF1 and IKZF3 for T-cells, CDX1 for stem cells, SOX30 and TCFL5 for male germ cells, and HEY1, ERG, and LHX6 for endothelial cells.

The ability to generate probabilistic associations based on learned patterns demonstrates TranscriptFormer’s internalization of complex gene regulatory networks during pretraining, enabling biologically meaningful predictions even for unseen data distributions. More importantly, this capability sketches a future path for using generative models as virtual instruments in place of cell atlases, whereby an integrated view of cells is not just a look-up table of data, but also a queryable interactive model of the knowledge contained that can simulate complex views into the data.

## Conclusions

TranscriptFormer represents a significant advancement in biological foundation models by successfully integrating transcriptomic data across an unprecedented evolutionary span of 1.53 billion years. Our findings demonstrate that generative pretraining on both genes and expression counts from diverse species creates representations with emergent biological properties that surpass previous approaches. The performance of TF-Metazoa and TF-Exemplar over single-species models on both in-distribution and out-of-distribution tasks provides compelling evidence that broader evolutionary pretraining enhances biological generalization. This suggests that evolutionarily conserved gene expression signatures can be effectively captured by foundation models, enabling robust cross-species analysis of cell types and states.

Beyond benchmarking, TranscriptFormer introduces a paradigm shift in how we interact with cellular data by functioning as a virtual instrument for biological inquiry. The model’s ability to generate meaningful cell type-specific transcription factor predictions and genegene interactions through prompting demonstrates its potential as a computational tool for hypothesis generation. Context-aware gene embeddings and cell embeddings further reveal how TranscriptFormer learns to encode multi-level biological structure—from cell types to tissues to donor-specific variations—without explicit supervision. This emergent capability points toward future models that could serve not merely as repositories of data but as interactive knowledge bases capable of simulating complex cellular phenomena.

Future work will focus on expanding the diversity of species represented, and mechanisms to better integrate diverse datasets by handling batch effects in a unified fashion, a limitation commonly shared across many foundation models. Similar to other single-cell foundation models trained on cell atlas datasets, TranscriptFormer is also not specialized for zero-shot perturbation prediction, which is a fruitful direction for future technical enhancements. Further, it will be exciting to work on incorporating additional modalities beyond transcriptomics, and developing more sophisticated prompting strategies to fully realize the potential of generative foundation models as virtual instruments for biological discovery. TranscriptFormer establishes a framework for this vision, demonstrating how machine learning can help unify our understanding of cellular diversity across evolutionary time.

## Author contributions

**Conceptualization**: James D Pearce, Stephen R Quake, Theofanis Karaletsos

**Data curation**: Sara E Simmonds, Benjamin Nelson

**Formal analysis**: James D Pearce, Sara E Simmonds, Gita Mahmoudabadi, Lakshmi Krishnan, Giovanni Palla, Ana-Maria Istrate, Alexander Tarashansky

**Methodology**:James D Pearce, Lakshmi Krishnan, Gita Mahmoudabadi, Giovanni Palla, Alexander Tarashansky, Theofanis Karaletsos

**Project administration**: Donghui Li, Stephen R Quake, Theofanis Karaletsos

**Software:** James D Pearce, Lakshmi Krishnan, Giovanni Palla, Alexander Tarashansky, Omar Valenzuela

**Visualization**: James D Pearce, Sara E Simmonds, Gita Mahmoudabadi

**Writing – original draft**: James D Pearce, Sara E Simmonds, Gita Mahmoudabadi, Ana-Maria Istrate, Theofanis Karaletsos, Giovanni Palla

**Writing – review & editing**: James D Pearce, Sara E Simmonds, Donghui Li, Stephen R Quake, Theofanis Karaletsos

## Data availability

Pretraining human and mouse datasets are available from [CZ CELLxGENE](https://cellxgene.cziscience.com/collections) (see Extended Data Table 1 for dataset IDs) and GEO (GSE247719).

Pretraining datasets are publicly available for *Caenorhabditis elegans* (GSE126954, GSE229022, GSE98561), *Danio rerio* (GSE202639, [Zebrahub](https://drive.google.com/drive/folders/1gB9u679gUqjX8P-pqjm6JtZL67eLoOGO)), *Drosophila melanogaster* ([Aging Fly Cell Atlas](https://hongjielilab.shinyapps.io/AFCA/), [Fly Cell Atlas](https://flycellatlas.org/), [Alzheimer’s Disease Fly Cell Atlas](https://hongjielilab.org/adfca/)), *Gallus gallus* (GSE181577), *Lytechinus variegatus* (GSE184538), *Oryctolagus cuniculus* ([Rabbit-Gastrulation2022](https://marionilab.github.io/RabbitGastrulation2022/#data)), *Plasmodium falciparum* (Malaria Cell Atlas), *Saccharomyces cerevisiae* (GSE125162), *Spongilla lacustris* ([GSE134912](https://www.ncbi.nlm.nih.gov/geo/query/acc.cgi?acc=GSE195790)), *Xenopus laevis* ([GSE195790](https://www.ncbi.nlm.nih.gov/geo/query/acc.cgi?acc=GSE134912)) (see Extended Data Table 2 for details).

### Evaluation datasets

*Danio rerio* (GSE234216), *Microcebus murinus* ([Tabula Microcebus](https://drive.google.com/drive/folders/1gB9u679gUqjX8P-pqjm6JtZL67eLoOGO)), *Petromyzon marinus* (E-MTAB-11087), *Stylophora pistillata* ([GSE166901](https://www.ncbi.nlm.nih.gov/geo/query/acc.cgi?acc=GSE234216)), *Xenopus tropicalis* (GSE113074).

### Spermatogenesis dataset

*Homo sapiens* E-MTAB-11063, *Pan troglodytes* E-MTAB- 11064, *Gorilla gorilla* E-MTAB-11065, *Macaca mulatta* E-MTAB-11068, *Callithrix jacchus* E-MTAB-11069, *Mus musculus* E-MTAB-11071, *Monodelphis domestica* E-MTAB-11072, *Ornithorhynchus anatinus* E-MTAB-11070, *Gallus gallus* E-MTAB-11073.

### Other evaluation datasets

Homo sapiens Tabula Sapiens 2.0 (subset of [Tabula Sapiens](https://cellxgene.cziscience.com/collections/e5f58829-1a66-40b5-a624-9046778e74f5)), COVID-19 human lung ([CZ CELLxGENE](https://cellxgene.cziscience.com/collections/2a9a17c9-1f61-4877-b384-b8cd5ffa4085)), Tahoe-100M ([huggingface](https://huggingface.co/datasets/tahoebio/Tahoe-100M)), Innate immune response dataset (E-MTAB-6773). See Extended Data Table 2 for details.

CELLxGENE pretrained embeddings from TF-Sapiens and TF-Exemplar are available for programmatic download on the CZ CELLxGENE portal for the [Census](https://cellxgene.cziscience.com/census-models).

## Code availability

The pretrained models are publicly available on [GitHub](https://github.com/czi-ai/transcriptformer) and on CZI’s [virtual cells platform](https://virtualcellmodels.cziscience.com/paper/transcriptformer-2025-04).

## Acknowledgments

We gratefully acknowledge the CZI AI infrastructure team for providing support with the GPU cluster required for training our models. We also want to thank the CZI Brand and Creative team for support in making and improving visual assets for this work. The CZI SciTech team was instrumental in sharing assets on our platform. We want to also acknowledge the scientific and technology leadership teams across the Chan Zuckerberg Science ecosystem including Andrea Califano, Sandra Schmid, Shana Kelley, Scott Fraser, Amy Herr, and Patricia Brennan for continuous feedback on this work, as well as Emma Lundberg, Yang Joon Kim, Mathais Voges and Jia Li Lim for their feedback and support.

## Materials and Methods

### Data

We curated an extensive collection of single-cell RNA sequencing datasets for model pretraining and evaluation from public sources comprising 115 million cells from 465 tissues, 1,865 cell types in 129 disease states across 23 species.

#### Pretraining Data Selection and Curation

For pretraining, we selected datasets that span the major animal lineages. This diverse dataset includes six vertebrate species (*Homo sapiens, Mus musculus, Oryctolagus cuniculus, Gallus gallus, Xenopus laevis, Danio rerio*), four invertebrate species (*Lytechinus variegatus, Caenorhabditis elegans, Drosophila melanogaster, Spongilla lacustris*), and two outgroups: a fungus (*Saccharomyces cerevisiae*) and a protist (*Plasmodium falciparum*). This phylogenetically rich collection spans approximately 1.53 billion years of evolutionary history [25] (Fig.1).

Most of these data were sourced from CZ CELLxGENE using the Discover API [49], comprising 57 million human cells (644 datasets) and 27 million mouse cells (88 datasets) (SI Table 1). We filtered these datasets to exclude spatial assays and duplicate cells. Both normal (67 millions cells) and diseased (17 million cells across 123 diseases) cells were retained.

To capture as many cell types and states as possible from other species, we selected whole-organism, developmental, and aging cell atlases for the remaining 16 datasets (SI Table 2). These were obtained from various public repositories and processed into a standardized h5ad format.

##### Processing

We converted each dataset to AnnData objects, verified the presence of raw gene count layers, harmonized metadata fields (`organism, assay, suspension type`), and mapped and updated gene features to Ensembl gene stable IDs (v113) for each species. Two datasets (*Spongilla lacustris*[1] and Zebrahub[50]) required reprocessing from raw sequencing reads, which we accomplished using Cell Ranger v9.1 with default settings and custom genome references (odSpoLacu1.1 and GRCz11, respectively). To ensure data quality, we applied minimal filtering to remove cells with fewer than 200 non-zero in-vocabulary genes, preserving the biological heterogeneity of the data.

#### Evaluation Data Selection and Curation

We assembled a comprehensive set of evaluation datasets to benchmark TranscriptFormer against existing methods.

##### Tabula Sapiens 2.0

A human scRNA-seq reference cell atlas [32]. We downloaded singlecell RNA sequencing data from the Tabula Sapiens Consortium via CZ CELLxGENE. To ensure independence from our training data, we filtered the “All Cells” dataset to include nine donor IDs absent from Tabula Sapiens 1.0 (TSP17 to TSP30). We excluded pancreas data due to insufficient cell counts (*>*100 cells).

##### Spermatogenesis dataset

A multi-species snRNA-seq dataset of testes from the major lineages of mammals and birds [27]. Data from nine species (*H. sapiens, Gorilla gorilla, G. gallus, Callithrix jacchus, Macaca mulatta, Monodelphis domestica, M. musculus, Ornithorhynchus anatinus, Pan troglodytes*) were downloaded from two sources (GEO and Bgee). Data were aligned and merged to create a dataset with raw counts (GEO) and harmonized metadata (Bgee).

##### Tabula Microcebus

A scRNA-seq reference cell atlas for mouse lemur (*Microcebus murinus*), a model primate species [51]. We obtained data from the Tabula Microcebus Consortium. We filtered out cells marked as low quality, contaminated, or lacking cell type assignments according to the original authors’ criteria (179,519 cells).

##### Tropical clawed frog cell atlas

A scRNA-seq developmental cell atlas for *X. tropicalis* covering embryonic stages [4]. The full embryonic dataset was subset to the early tailbud stage when dozens of cell types have differentiated (34,335 cells; 54 cell types).

##### Zebrafish cell landscape

A scRNA-seq developmental cell atlas for *D. rerio* [52]. The dataset was subset to adult stage and split by tissue (N=10) (184,808 cells; 91 cell types). All embryonic stages were pooled.

##### Stony coral cell atlas

A scRNA-seq reference cell atlas for stony coral (*Stylophora pistillata*) [53]. The dataset was subset to include adult stage only (15,252 cells; 13 cell types).

##### Sea lamprey brain

A snRNA-seq dataset of brain tissue from *Petromyzon marinus* [54]. This dataset required reprocessing of the raw sequencing reads. We used Cell Ranger v9.1 with default settings and a custom genome reference (Pmarinus7.0).

##### COVID-19 human lung

The scRNA-seq experimental dataset of four healthy donors’ lung sections infected with SARS-CoV-2 [33]. Data were downloaded from CZ CELLxGENE.

##### Tahoe-100M

A comprehensive human scRNA-seq drug perturbation dataset comprising 100 million cells from cell lines [24]. We downsampled the dataset to 100,000 cells from plate 3, resulting in 95 unique small molecule perturbations over 50 cell lines.

##### Innate immune response dataset

A multi-species single-cell RNA sequencing dataset characterizing innate immune responses in fibroblasts and mononuclear phagocytes challenged with LPS perturbations in mouse, rat, rabbit, and pig [28].

##### Processing

We converted all datasets to AnnData objects, harmonized metadata fields (`organism, cell type`), and mapped gene features to Ensembl gene stable IDs (v113). For quality control we removed duplicate cell barcodes. For all cell type prediction tasks, we applied consistent curation steps across evaluation datasets. We removed cells with ambiguous cell type annotations (labeled as “cell,” “unassigned,” “unknown,” or “Unclassified”) and filtered out genes with zero counts across all cells. We also standardized all count matrices to be raw counts, ensuring consistency across all datasets for our analyses.

### Modeling Cells as Bags of Transcripts

Cells can be represented as *bags of transcripts*, where each transcript *t*_*k*_ corresponds to an observed mRNA molecule. These transcripts are unordered, and identical transcripts are indistinguishable from one another, meaning that the representation of a cell is based on the *multiset* of observed transcripts rather than their order.

Mathematically, let a cell be represented as a collection of transcripts:

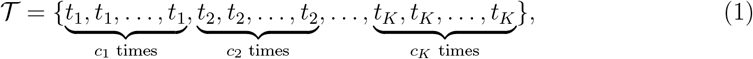

where *K* is the number of unique transcripts observed. Because transcripts are sampled from expressed genes, the same transcript can appear multiple times within a cell. By aggregating transcripts at the gene level, we define a *gene expression profile* instead as a collection of genes, where each gene *g*_*j*_ is associated with a *count c*_*j*_, denoting the number of times a transcript from that gene was observed.

Formally, the gene expression profile, 𝒢, representation of a cell is:

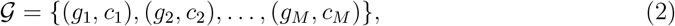

where *g*_*j*_ is a unique gene with non-zero counts; *c*_*j*_ *>* 0 is the *expression level* of gene *g*_*j*_, defined as the number of times a transcript from gene *g*_*j*_ appears; *M* is the number of expressed genes in the cell.

This transformation from transcripts to gene-level counts allows for a more efficient representation of the bag of transcripts, capturing the overall expression profile rather than individual transcript occurrences. The next section describes a generative model that formalizes this process, modeling the sequential selection of genes and their associated expression levels.

### A Generative Model for Gene Expression

The generative model for gene expression represents the process by which a cell’s transcriptomic profile is formed. Given a set of genes and their associated expression levels, we model the generation of a cell’s expression profile as a sequential process involving two key steps: (1) selecting a gene from a categorical distribution and (2) sampling its expression count from a zero-truncated Poisson (ZTP) distribution.

#### Step 1: Gene Selection

At each step *j*, a gene *g*_*j*_ is sampled from a categorical distribution over the gene vocabulary. The probability of selecting a gene depends on previously chosen genes and their counts. Formally, the probability of selecting *g*_*j*_ at step *j* is given by:

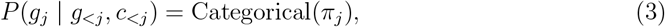

where *π*_*j*_ = *f*_*θ*_(*g*_*<j*_, *c*_*<j*_) is a probability distribution over the gene vocabulary, parameterized by the function *f*_*θ*_ with learned parameters *θ*.

#### Step 2: Count Sampling

Once a gene *g*_*j*_ is selected at step *j*, its expression level *c*_*j*_ is sampled from a zero-truncated Poisson (ZTP) distribution. The ZTP distribution ensures that every selected gene has a strictly positive count:

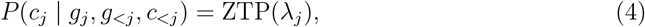

where *λ*_*j*_ = *f*_*ϕ*_(*g*_*j*_, *g*_*<j*_, *c*_*<j*_) is the Poisson rate parameter, estimated by the function *f*_*ϕ*_ with learned parameters *ϕ*.

#### Joint Distribution over Genes and Counts

The generative process described above defines a structured distribution over both the genes present in a cell and their corresponding expression levels. We further introduce the assay type, *a*, which is condtion on. Given the sequential nature of the model, the joint probability distribution over all genes and counts in a cell is factorized as:

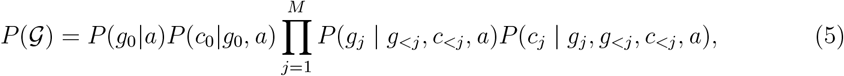

Since the categorical distribution for gene selection and the ZTP distribution for count sampling are both conditioned on the previously sampled genes and counts, this formulation captures dependencies between genes in a cell’s expression profile. The parameters *θ* and *ϕ* are jointly learned, allowing the model to infer gene co-expression patterns and contextdependent count distributions.

This generative framework provides a structured probabilistic model for single-cell transcriptomics, enabling explicit modeling of gene-gene interactions and expression dependencies within a cell capturing the structured nature of transcriptomic data.

### Model Architecture

#### Input Embeddings

TranscriptFormer embeds gene sequences across diverse species and sequencing technology into continuous representations. For each gene *g*_*j*_, we map it to its associated amino acid and proteins sequences. These sequences are embedded using ESM-2 [15], a protein language model. If there are multiple proteins associated with a gene (e.g., protein isoforms) then the mean over all ESM-2 protein embeddings is used.

Additionally, we introduce an *assay token* **a** ∈ ℝ^*d*^ that encodes information about the sequencing technology used. The transformer input, *Z*^(0)^ ∈ ℝ^(*M* +1)×*d*^, combines the gene embeddings (processed through an MLP) with the assay token:

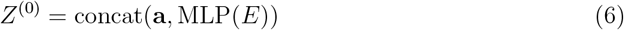

where 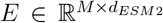 is the protein embeddings tensor. The MLP serves to project the ESM-2 embedding dimension *d*_*ESM*2_ to the model dimension *d*.

#### Transformer Architecture

The model follows a standard transformer encoder architecture with *L* stacked layers. The output of the *l*-th layer is defined recursively as:

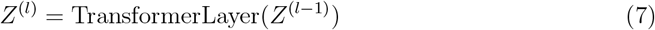

where *Z*^(*l*−1)^ ∈ ℝ ^(*M* +1)×*d*^ is the output from the previous layer. We employ pre-normalization, GeLU activation as well as removal of MLP and attention bias terms. We omit positional encodings from the input since the gene expression profiles have no inherit sequence.

#### Expression-Aware Multi-Head Self-Attention

We encode count information as a bias term in the attention matrix at each transformer layer. We modified the attention mechanism to incorporate the counts as follows:

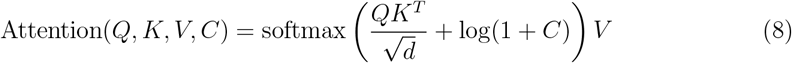

where the count matrix *C* ∈ ℝ^(*M* +1)×(*M* +1)^ is constructed by repeating the count vector **c** = (1, *c*_1_, *c*_2_, …, *c*_*M*_) across all rows. *Q, K* and *V* are the standard query, key and value matrices respectively [55]. In Supplementary Methods we show that this bias term ensures that higher-count genes exert greater influence in the attention computation in a way that is consistent with the bag-of-transcripts representation of cells. For numerical stability, we apply soft capping to the softmax logits. The standard causal attention mask is applied across gene tokens, while the assay token is never masked.

#### Decoders Heads

The gene decoder head predicts the next gene categorical distribution parameters *π*_*j*+1_ at each position *j* using a two-layer MLP followed by softmax:

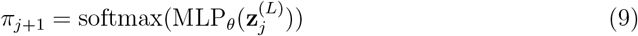

where 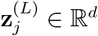 is the *j*th sequence component of *Z*^(*L*)^.

The input to the count decoder is obtained by concatenating the final hidden states with the transformer input for the next position *j* + 1, which is then passed through a MLP to get the pre-normalized count logits **u** ∈ ℝ^*M*^ as so

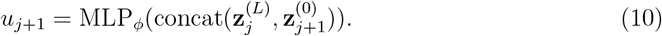

To improve training stability, a softmax normalization is applied to **u** (across the sequence dimension) and then rescaled to ensure that the predicted counts sum to the observed total *N* = ∑*c*_*j*_. Additionally a learnable gene-level bias **b** ∈ ℝ^|*V* |^ is applied to all genes in the input sequence to obtain the Poisson rate parameter vector:

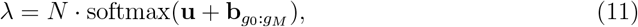

where **b**_*g*0:*gM*_ denotes the vector of bias terms for non-zero genes.

#### Cell and Contextual Gene Embedding

TranscriptFormer produces two types of embeddings: cell embeddings and contextual gene embeddings (CGEs). Cell embeddings represent the entire expression profile of a cell as a single vector, computed as the average over all transformer outputs (excluding the assay token) from the final layer:

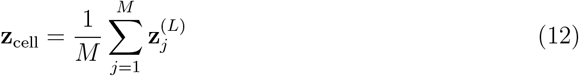

where 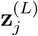 denotes the output of the *L*-th transformer layer for the *j*-th gene position. Contextual gene embeddings (CGEs), on the other hand, capture gene-specific representations within their cellular context. Each CGE corresponds directly to an individual component 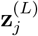 of the transformer’s final layer output.

#### Loss Functions

The model is trained using two distinct loss functions corresponding to the two decoder heads: a cross-entropy loss for gene prediction and a zero-truncated Poisson negative loglikelihood (NLL) loss for count prediction.

#### Gene Decoder Loss

The gene decoder predicts a probability distribution over the gene vocabulary at each sequence position. The standard cross-entropy loss is used to supervise the gene predictions. Given the predicted gene probabilities *π*_*j*_ at position *j* the gene loss is given by:

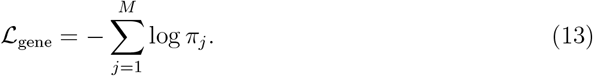

This loss encourages the model to assign high probability to the correct gene identity at each position.

#### Count Decoder Loss

The count decoder predicts a set of expected expression counts per sequence position. Since gene expression counts follow a discrete, non-negative distribution, we model them using a Poisson likelihood. However, because counts of zero are excluded from the input sequence, we employ a zero-truncated Poisson negative log-likelihood (NLL) loss.

Given the predicted rate parameter *λ*_*j*_ for position *j* and the observed ground truth count *c*_*j*_, the loss is computed as:

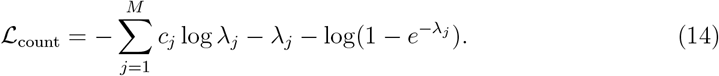

The additional term log(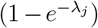) accounts for the truncation of zero values, ensuring proper normalization.

#### Total Loss

The final loss function is the sum over gene and count losses:

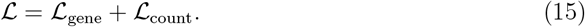

This formulation ensures that the model jointly learns to predict gene identities and their corresponding expression levels in a biologically meaningful manner.

### Model Configurations and Versions

We trained three variants of the TranscriptFormer architecture, each using a different subset of the available dataset, but all sharing an identical core model configuration. Each model is a 12-layer transformer with 16 attention heads per layer, using a model dimension and hidden size of 2048 and GELU activation. The architecture employs full-sequence selfattention with a maximum sequence length of 2,047 tokens (representing genes), no forward or attention biases, and a dropout rate of 0.1 applied throughout. The gene embeddings are initialized from pretrained representations and held fixed during training. This consistent architecture results in a transformer module with approximately 302 million parameters, with overall trainable parameters varying across models due to differences in vocabulary size and decoder head dimensionality.

To handle the exceptionally large output vocabulary in the TF-Metazoa model (over 247,000 genes across 12 species), we reduced the output embedding dimensionality from 2048 to 512, which yielded no loss in performance during preliminary evaluations.

Following the TranscriptFormer descriptions, we include the benchmark models used for comparison.

**TF-Metazoa** was trained on 112 million cells spanning all twelve species described in the Data section. This dataset includes six vertebrate species (*Homo sapiens, Mus musculus, Oryctolagus cuniculus, Gallus gallus, Xenopus laevis, Danio rerio*), four invertebrate species (*Lytechinus variegatus, Caenorhabditis elegans, Drosophila melanogaster, Spongilla lacustris*), and two outgroups: a fungus (*Saccharomyces cerevisiae*) and a protist (*Plasmodium falciparum*). The model includes 444 million trainable parameters and 633 million non-trainable parameters (from frozen pretrained embeddings). This model has a total vocabulary size of 247,388.

**TF-Exemplar** was trained on 110 million cells from five model organisms: human (*H. sapiens*), mouse (*M. musculus*), zebrafish (*D. rerio*), fruit fly (*D. melanogaster*), and *C. elegans*. Total trainable parameters: 542 million; non-trainable: 282 million. This model has a total vocabulary size of 110,290.

**TF-Sapiens** was trained on 57 million human cells. This model has 368 million trainable parameters and 61 million non-trainable parameters. This model has a total vocabulary size of 23,829.

**UCE (Universal Cell Embedding)** is a multi-species foundation model trained on an integrated atlas of about 36 million single cells spanning eight species (human, mouse, mouse lemur, zebrafish, pig, rhesus macaque, crab-eating macaque, and western clawed frog) [22]. The model uses a 33-layer transformer with approximately 650 million parameters. Notably, UCE represents each gene via a pretrained protein embedding (from the 15B-parameter ESM2 model) to incorporate protein-sequence information. During pretraining, a portion of expressed genes in a cell are masked and the model is optimized to predict whether a given gene was expressed, using the cell’s embedding and gene protein-token embeddings as context.

**scGPT** is a transformer-based model with approximately 53 million parameters, pretrained on around 30 million human single-cell transcriptomes [18]. Continuous gene expression values are discretized into categorical bins, and the model uses a BERT-style masked language objective to predict masked gene expression bins from context genes.

**scVI (CellxGene version)** is a variational autoencoder-based deep generative model [31]. For our benchmarks the scVI model was trained on non-spatial, human RNA sequencing data from the CZ CELLxGENE Discover Census, version 2024-07-01 (74.3M cells). Although it has fewer parameters (7.1 million), it serves as a strong baseline due to its robust probabilistic modeling.

**AIDO.Cell-100M** is an encoder-only transformer model with 100 million parameters trained on 50 million single human cells that span more than 100 tissue types [29]. It employs an auto-discretization of gene expression values and uses a read depth-aware denoising training objective, teaching the model to recover original expression counts from artificially subsampled transcriptomes.

**Geneformer (12L-30M-i2048)** is a context-aware transformer model trained on approximately 30 million human single-cell transcriptomes (Genecorpus-30M) [30]. Each cell is represented by genes ranked by expression levels, and the model predicts masked genes from their context, capturing gene network interactions. The benchmark variant used here has 12 layers, 8 attention heads, hidden size of 512 and max sequence length of 2048 input genes per cell.

**ESM2-CE (ESM-2 Cell Emebdding)** is our own benchmark model based on the ESM2 model [15]. This model consists of the ESM-2 embedding inputs to TranscriptFormer, vectored averaged over the sequence dimension. It serves as a baseline for assessing the utility of TranscriptFormer and single-cell pre-training over the ESM-2 model.

### Model Training

#### Training Setup

Training was conducted on CZI’s compute cluster equipped with 1000 H100 GPUs, using 16-bit mixed precision for improved efficiency and memory utilization. The training loop was implemented using the the PyTorch Lightning with Distributed Data Parallel (DDP). We used PyTorch’s Flex Attention [56] to build the transformer blocks. A global batch size of approximately 4–5 million tokens was used, achieved with a per-GPU batch size of 4–16 examples and gradient accumulation steps of 6. Gradients were clipped to a maximum norm of 1.0 to improve stability.

#### Training Data Sampling

During training, expression profiles were sampled from each species *s* with sampling probability *p*_*s*_, where the sampling probability is given by:

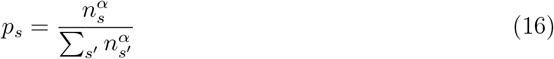

where *n*_*s*_ is the number of training cells from species *s*, and *α* = 0.6 controls the sampling skew. This strategy increases the relative sampling frequency of low-resource species while preventing overfitting to high-resource species like human or mouse.

#### Sequence Processing

We trained with a sequence length of 2048, which includes the gene expression profiles and assay token. Genes were randomly shuffled in each batch to enforce permutation invariance. Sequences longer than 2048 were truncated, and shorter sequences were padded using a special [PAD] token. The model was trained using causal masking over this gene vocabulary, with raw count values clipped to a maximum of 30. Cells with less than 200 expressed gene were filtered out of training. Additionally, outlier cells with expression levels exceeding 3 standard deviations above or below the dataset average were filtered.

#### Optimization

We used the AdamW [57] optimizer with *β*_1_ = 0.9, *β*_2_ = 0.95, and weight decay set to 0.1. The learning rate was scheduled using a linear warm-up over the first 10% of training steps, followed by cosine decay to the minimum learning rate. The peak learning rate was set to 1.5 × 10^−4^, decaying to a minimum of 1.0 × 10^−5^. Learning rates were scaled relative to a baseline batch size of 1,536 tokens to maintain consistency across model variants. Each model was trained for a total of approximately 3.5 trillion tokens, equivalent to 15 epochs for the TF-Metazoa model.

### Analysis

#### Linear Probing for cell type and cell state classification

Linear probing is a technique used to evaluate the quality of learned representations by training a simple linear classifier on top of fixed, pre-trained embeddings. This approach assesses how well the underlying representations capture relevant information for downstream tasks without fine-tuning the entire model.

Formally, given a pre-trained model *f*_*θ*_ that maps input data *x* to embeddings *e* = *f*_*θ*_(*x*), linear probing involves training a linear classifier *g*_*w*_ with parameters *w* to predict target labels *y* from these fixed embeddings:

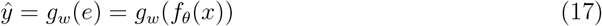

The parameters *θ* of the pre-trained model remain frozen during this process, while only the parameters *w* of the linear classifier are optimized.

When evaluating performance of the TranscriptFormer models, as well as the benchmark models, we use Logistic Regression (LR) with L2 regularization penalty (*C* = 0.01) implemented in scikit-learn [58]. We used 5-fold cross-validation to estimate the performance of the models. We report the macro F1 score for each cell type and averaged across all cell types. We also report the standard error of the F1 score across the 5 folds.

#### Cross-Species reference mapping for cell type and cell state transfer

To assess the ability of TranscriptFormer to generalize transcriptional representations across species, we performed a cross-species reference mapping experiment using a k-nearest neighbors (kNN) classifier.

We adopted a transfer learning framework in which labels from one species (source species) are transferred and evaluated on embeddings from a different species (target species). For each source-target species pair, we embedded all cells using TranscriptFormer or one of the benchmark models and extracted their representation from the final encoder layer. A kNN classifier implemented in scikit-learn [58] (*k* = 10) was use with the source species’ cell embeddings and labels as reference, which are mapped onto the target species to predict its labels with a majority vote.

Classification performance was quantified using the F1 score, calculated per cell type and averaged to obtain a macro F1 score (see Supplementary Methods) for each source–target species pair. This reference mapping approach evaluates a model’s ability to transfer cell type and cell state information without fine-tuning or training of an auxiliary model.

#### COVID-19 disease classification task

For the disease classification task, we downloaded the dataset from CELLxGENE from the corresponding publication Wu et al. [33], see. We used the original labels of “infected” cells (True or False) provided by the original authors. We adopted the linear probing evaluation discussed above.

#### Contextualized Gene Embedding

To systematically evaluate contextualized gene embeddings (CGEs), we developed several types of CGE analyses. These included cross-donor, cross-cell type, and cross-tissue comparisons, as well as a flexible variance partitioning framework for quantifying the influence of different biological covariates.

##### Gene Selection

For each analysis type, raw count matrices from Tabula Sapiens 2.0 datasets were subset to the relevant cell types (typically those containing large numbers of cells) and preprocessed using Scanpy [59]. Standard preprocessing steps were applied, which included gene filtering (minimum of 3 cells), library-size normalization to 10,000 counts per cell, and log-transformation. Highly variable genes (HVGs) were identified using the Seurat method [10]. In cross-tissue and cross-donor analyses, genes were retained if they were expressed in at least 10% of cells within the target cell type. For cross-cell type comparisons, genes were required to exceed the same threshold in at least three distinct cell types. This was done to increase the probability that the selected genes would have a corresponding CGE in a large number of cells, as the model selects only the top 2047 expressed genes per cell to form each cell sentence. A fixed number of valid genes (*n* = 20) were randomly selected per analysis, and those without a corresponding CGE in any cells were excluded from the analysis. We created an auxiliary function for retrieving additional metadata (gene functional annotation) for the selected genes based on their Ensembl IDs.

##### Embedding Aggregation and Dimensionality Reduction

CGEs were obtained from inference runs through the TranscriptFormer model. Cells were grouped by donor (cross-donor), tissue (cross-tissue), or cell type (cross-cell type), and gene embeddings were averaged within each group. This process also recorded the number of cells contributing to each averaged CGE.

The averaged CGEs were projected into two dimensions using principal component analysis (PCA). PCA results were visualized in multi-panel scatter plots: (1) colored by the biological grouping variable (e.g., donor, tissue, or cell type), (2) colored by gene identity, and (3) optionally colored by the log_2_-transformed number of contributing cells. These plots allowed direct visual comparison of the embedding structure across biological axes.

##### Variance Partitioning Analysis and Gene Selection Tests

To assess the impact of gene selection criteria on the observed CGE trends, we implemented a flexible analysis that supports parameterization of gene filtering based on HVG status, expression thresholds, and the minimum number of cell types in which a gene must be expressed. For each configuration, the CGEs were aggregated by donor, tissue, and cell type, followed by PCA.

We evaluated four gene selection criteria (see Supplementary Methods for additional details on the selected criteria):

- **Baseline** HVG filtering enabled, expression threshold = 10%, min cell types = 3
- **Test 1** HVG filtering disabled, expression threshold = 10%, min cell types = 3
- **Test 2** HVG filtering disabled, expression threshold = 1%, min cell types = 3
- **Test 3** HVG filtering disabled, expression threshold = 1%, min cell types = 1

Each configuration was run using the same number of randomly selected genes. For each result, variance partitioning [60] was performed by fitting linear models to PCA coordinates and computing partial *R*^2^ values for donor, tissue, and cell type. The results are shown in SI Fig. 7.

#### Pairwise Gene-Gene Interaction Analysis

To assess whether TranscriptFormer captures biologically meaningful regulatory relationships between transcription factors (TFs) and their target genes, we computed pointwise mutual information (PMI) [61] values for each TF–gene pair, conditioned on a specified assay token *a*:

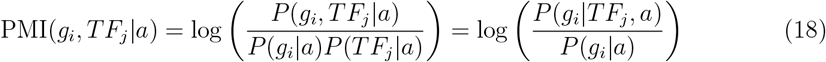

Marginal probabilities *P* (*g*_*i*_|*a*) were obtained via single-step decoding from TF-Sapiens by “prompting” on the assay token, while conditional probabilities *P* (*g*_*i*_|*TF*_*j*_, *a*) were derived from two-step decoding, i.e., prompting with *TF*_*j*_ as the initial token (*c*_*j*_ = 1). To reduce spurious effects from low-probability genes, we filtered out genes with marginal probabilities below 10^−8^, retaining 17,625 protein-coding genes for analysis.

To assess statistical significance of interactions, we computed Z-scores for each PMI value:

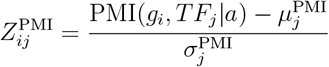

where 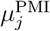 and 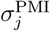 denote the mean and standard deviation, respectively, of the PMI distribution for transcription factor *T F*_*j*_, estimated across all considered genes. This per-TF background modeling accounts for variability in interaction distributions between different transcription factors. Two-sided p-values were then calculated using the standard normal distribution as:

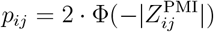

where Φ is the standard normal cumulative distribution function. Resulting p-values were adjusted for multiple hypothesis testing using the Benjamini–Hochberg procedure [62] to control the false discovery rate (FDR). We applied an FDR threshold of *α* = 0.05 to identify statistically significant TF–target gene interactions inferred by the model.

#### Cell Type-Specific Transcription Factor Generation

To assess the ability of TranscriptFormer to capture cell type-specific transcription factor (TF) activity, we implemented a conditional generation strategy inspired by the Tabula Sapiens 2.0 (TS2) study [32]. We began by identifying the top 100 marker genes for each of 112 cell types spanning 28 tissues in the TS2 dataset. For a given cell type, *s*, the set of marker genes ℳ (*s*) were selected based on average expression profiles within each cell type, resulting in a high-confidence gene set for conditioning the model.

Using these cell type-specific gene signatures, we probed the TF-Sapiens model by prompting it with the set ℳ (*s*) alongside an assay token *a*, and performed a single forward decoding step to obtain the conditional log-probability distribution over all 1,635 human transcription factors in the TS2 study:

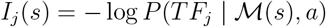

where we interpret *I*_*j*_ (*s*) as the surprise of observing the activation of transcription factor *T F*_*j*_ in a cell of type *s*. This probabilistic output reflects the model’s internal representation of TF–cell type associations, even though it was not explicitly trained on the TS2 dataset.

To identify dominant TFs for each cell type, we grouped transcription factors with the cell type that minimizes the surprise of observing it. That is, for each transcription factor *T F*_*j*_, we identified the most associated cell type *s* ^∗^ as:

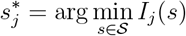

𝒮where is the set of all 112 cell types. This allowed us to map transcription factors to the most likely associated cell type according to the model’s learned internal representations.

The resulting matrix of (negative) surprise values across all TF–cell type combinations was visualized as a heatmap (Fig.5B). Transcription factors were arranged according to their assigned cell type, revealing distinctive structural features: vertical bands corresponding to ubiquitously expressed TFs and diagonal patterns highlighting cell-specific regulatory factors.

#### Cell type centroid and similarity computation

The axes of biological variation in cellular gene expression patterns are both across (e.g., phylogenetic) and within species (e.g., differentiated cell types, developmental cell states, tissue-specific context). We studied whether TranscriptFormer cell embeddings captured multiple levels of biological variation by comparing cell types and states within and between species.

To enable robust cross-species and cell type comparisons, we computed cell type centroids from TranscriptFormer cell embedding vectors. For each species and cell type combination, individual cell embeddings were grouped by author provided cell type label. The cell type centroid was calculated as the mean across all cells of that type within each species. This approach reduces technical and biological noise inherent in individual cell measurements while preserving the core transcriptional identity of each cell type, yielding stable representative vectors suitable for downstream analyses. To quantify transcriptional relationships across all cell types and species, we computed pairwise similarity matrices between cell type centroids. For each pair of species, cosine similarities were calculated between all possible cell type centroid combinations using SciPy cdist function with the cosine distance metric.

#### Within species - Differentiation trajectories

To test whether TranscriptFormer embeddings learned the developmental trajectory of cells during spermatogenesis, we attempted to reconstruct the known developmental sequence from cell type similarity patterns using human data from the Spermatogenesis dataset (Murat et al., 2023). Per cell type cosine similarities were calculated from the TF-Metazoa embedding to create a pairwise similarity matrix. The known biological developmental sequence (male germ line stem cell, spermatogonium, spermatocyte, early spermatid, late spermatid) was used as ground truth to evaluate reconstruction accuracy.

#### Across species - Phylogenetic structure

We examined whether TranscriptFormer cell embeddings capture phylogenetic structure in gene expression patterns. If the embeddings capture phylogenetic structure then we would expect closely related species to have more similar gene expression patterns than distantly related species for shared cell types. We conducted an evolutionary analysis using data from eight species in the Spermatogenesis dataset, comparing TF-Metazoa and ESM2-CE embeddings as the baseline. For each species, we created pseudo-bulk embedding centroids by averaging embeddings across cells, using only the five cell types (early spermatid, late spermatid, somatic cell, spermatocyte, spermatogonium) common to all species to ensure consistent comparison. We then calculated pairwise cosine distances between species vectors. To quantify how well these embedding distances reflect evolutionary relationships, we compared them with established evolutionary distances (in millions of years) from TimeTree [25] using Spearman’s rank correlation.

#### Cell type similarity search across species

To test whether TranscriptFormer cell embeddings could be used to map cell type similarities across species to study cell type evolution, we constructed a 1.6M cell developmental atlas spanning 5 phylogenetically diverse species representing major evolutionary lineages of animals. The atlas included an invertebrate (round worm (*C. elegans*) [63] and vertebrates (zebrafish (*D. rerio*) [64](Saunders et al., 2023), clawed frog (*X. tropicalis*) [4], rabbit (*O. cuniculus*) [65], pig (*S. scrofa*) (Simpson et al., 2024) and each species dataset encompassed multiple developmental stages (see Methods). This comprehensive atlas enabled systematic comparison of developmental programs across phylogenetically distant taxa, providing the foundation for evaluating whether learned embeddings preserve evolutionary and developmental relationships in high-dimensional representation space. We compared species across varying phylogenetic distances: within mammals (rabbit/pig 86 MYA), across vertebrates (clawed frog/zebrafish 429 MYA; rabbit/zebrafish 429 MYA) and across the protostome-deuterostome split (clawed frog/worm 600 MYA).

Cell type centroids were computed as described above for each species by averaging embeddings within author provided cell type labels, requiring a minimum of 40 cells per cell type. Cross-species cell type matches were identified using similarity search, where similar cell types were detected by computing cosine similarities between all pairwise combinations of cell type centroids across species, only top matches were considered. Cross-species cell type relationships were visualized using interactive Sankey diagrams. Cell types from each species were positioned as source nodes (left side) and target nodes (right side), with flow connections representing similarity relationships above a specified threshold.

To assess whether cell type mappings were consistent with known biology, we examined each mapping manually. Matched cell types were considered biologically consistent when cell type labels were equivalent (e.g., Primordial Germ Cell (PGC), primordial germ cell), synonyms of each other (e.g., somite, muscle cell; blood progenitor, megakaryocyte–erythroid progenitors; gut/hypoblast, gut tube) or originating from analogous anatomical structures within established developmental hierarchies (e.g., Hindbrain, differentiating neuron (hindbrain), neural progenitor (hindbrain)). Cell type labels that did not meet this criteria, especially across distantly related species, were considered inconsistent, but potentially biologically analogous based on similarity of gene expression patterns (e.g., frog cement gland, worm pharyngeal gland).

#### Cell state similarity search across kingdoms

To explore whether TranscriptFormer cell embeddings can be used to search for similar cell states across unicellular fungi (yeast) and multicellular organisms, we compared yeast cells grown in ten different environmental conditions [66] to the animal developmental cell atlas. We used the same methods as described above, computing the cell type/state centroid per species, and calculating the cosine similarity between all cell centroid-to-centroid pairs for all species and recording the top matches for each yeast cell state.

## Supplementary Text

### Contextulized gene embedding analysis

#### Variance partitioning quantifies the contribution of cell type, tissue and donor information to Contextualized Gene Embeddings

To further quantify the contribution of each of the biological metadata (e.g. cell type, tissue and donor) to CGEs total “information content”, we first performed PCA on the CGEs derived from three tissues, three donors, and several cell types (SI Figure 6). We then employed a variance partitioning approach using nested linear regression models to quantify the unique contribution of three categorical predictors—cell type, tissue, and donor—to the variance in the first two principal components. The full model including all predictors was compared to reduced models that omitted one predictor at a time. Partial R^2^ values were calculated as the relative increase in the residual sum of squares (SSE) when a predictor was excluded from the full model. For PC1 and PC2, cell type accounted for *>* 95% of the explained variance when controlling for the other two variables (SI Figure 7). These findings agree with the clustering patterns clearly depicted in SI Figure 6, and demonstrate that the primary source of variability in the CGE space is cell type, with tissue and donor identity exerting only modest secondary effects.

**Figure 6.**
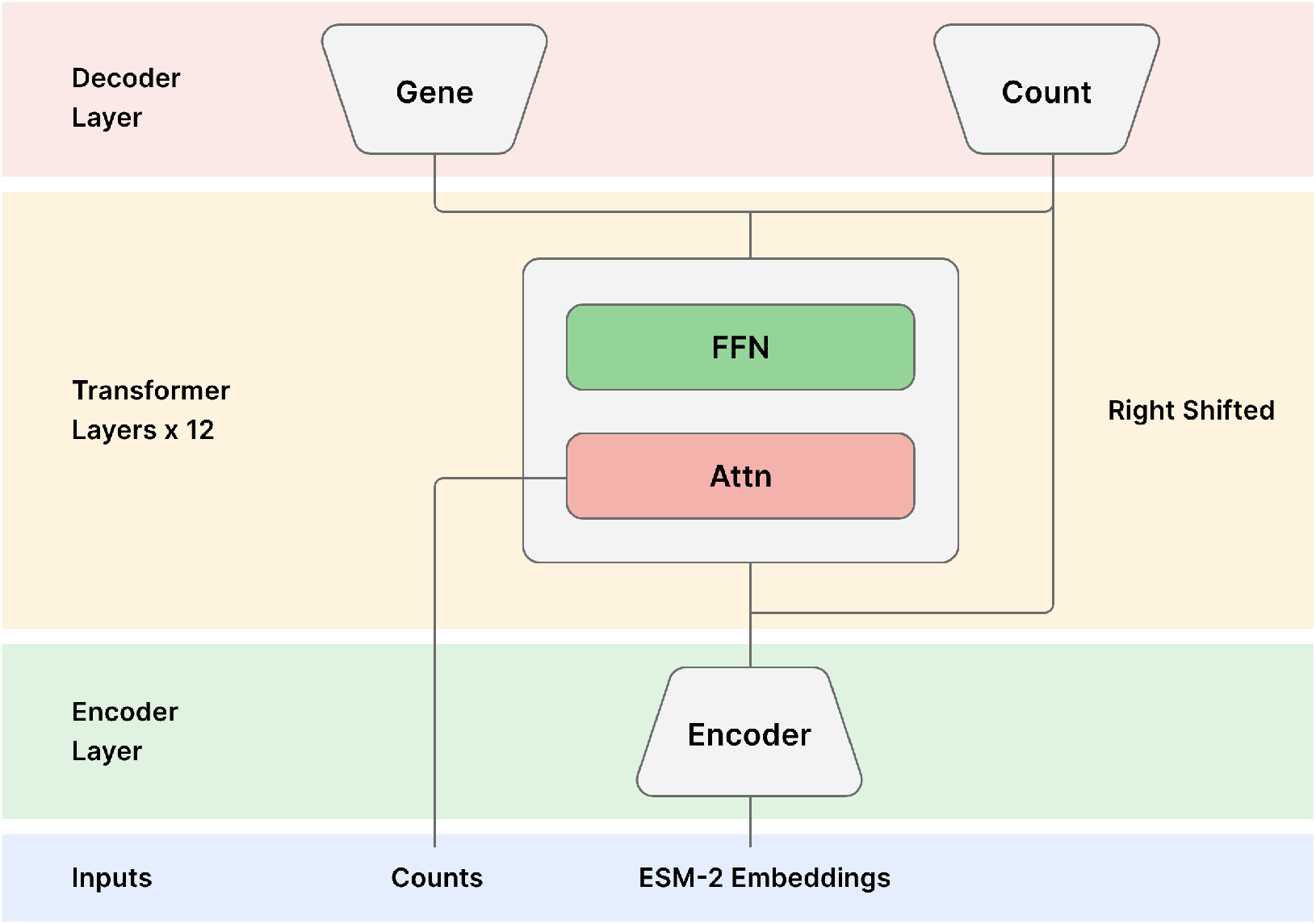
Model architecture diagram for TranscriptFormer models.

#### Observed trends are robust to changes in gene selection methodology

Additionally, we tested the impact of several gene filtering thresholds on the resulting PCA plots. The genes are selected from highly variable genes that appear across half of the cell types in the analysis and that are expressed across 10% of cells in each of those cell types (see Materials and Methods: CGE). To explore whether selection from highly variable genes has an influence on the output CGE PCA clustering patterns, we removed that filter, while keeping other filters intact. We obtained similar *R*^2^ values (denoted as “Test 1” in SI Figure 7) to those shown previously (denoted as “baseline” in SI Figure 7). Next, we additionally lowered another filter threshold for the percentage of cells within each type that are required to express a given gene, lowering the value from 10% to 1% (denoted as “Test 2” in SI Figure 7). Lastly, in addition to modifying these two filters, we removed the filter where at least half of the cell types in the analysis were required to express the final selected genes (denoted as “Test 3” in SI Figure 7). The resulting *R*^2^ values consistently show that the CGE PCA clusters are fairly robust to changes in filters applied to select genes, and that the primary context contained in CGEs is cell type, with tissue and donor contexts contributing less to the overall information content of CGEs. However, it appears that as some filters are relaxed, tissue context becomes slightly more predictive than donor context for PC2.

#### Exploring the impact of batches on CGE donor separation

We further constructed CGE tables based on batches for blood neutrophils from Tabula Sapiens 2 (SI Figure 8). Based on the clustering pattern, the overall observed trend is that CGEs from the same donor appear to cluster together and within each donor-cluster batches form subclusters. This suggests that donor effects are stronger than batch effects, however, we cannot completely rule out batch effects from donor effects.

## Supplementary Figures

**Fig. S1:**
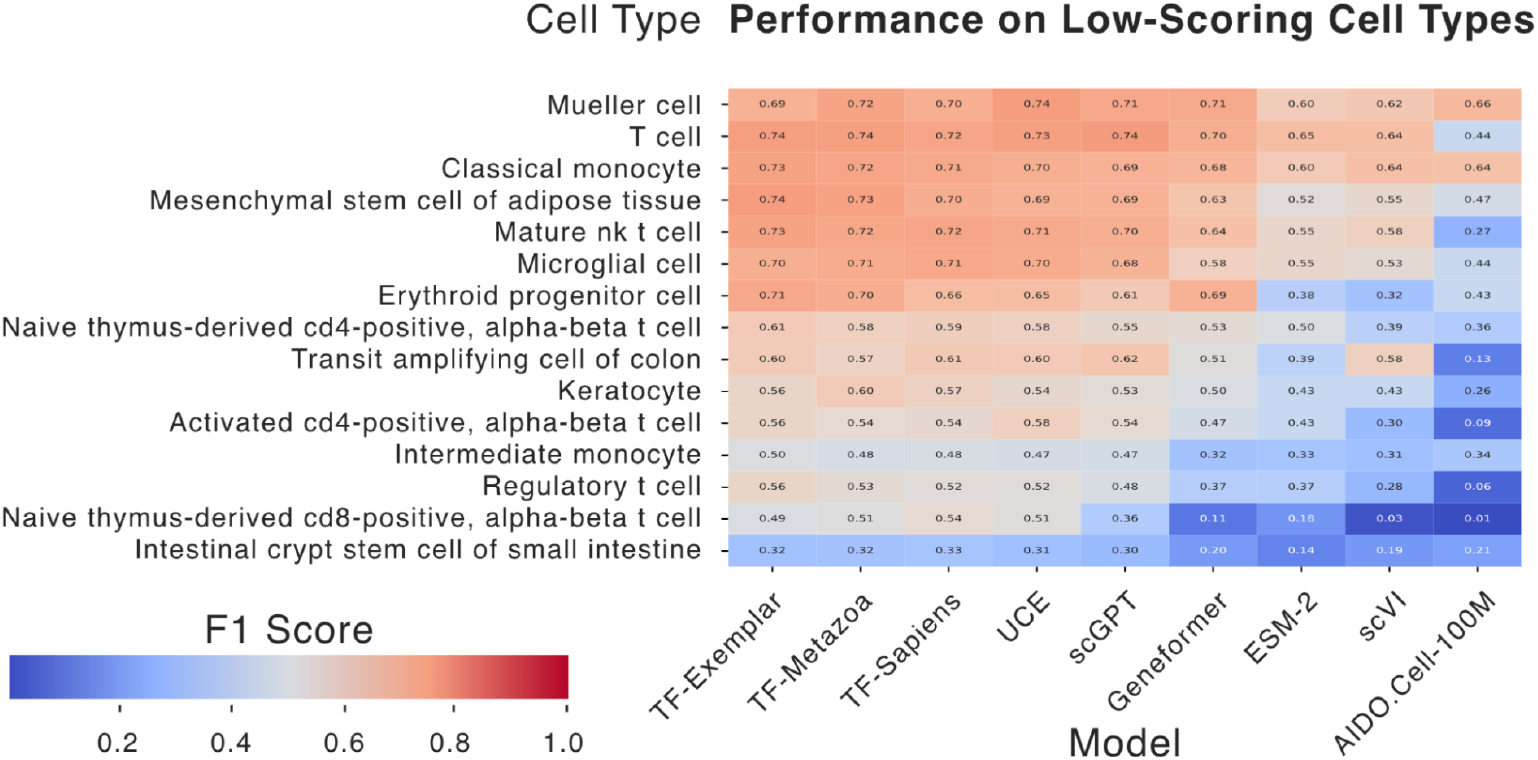
Tabula Sapiens 2.0 cell-type classification results for fer-cell-type F1 scores across models for select cell types that have the lowest mode-average F1 scores.

**Fig. S2:**
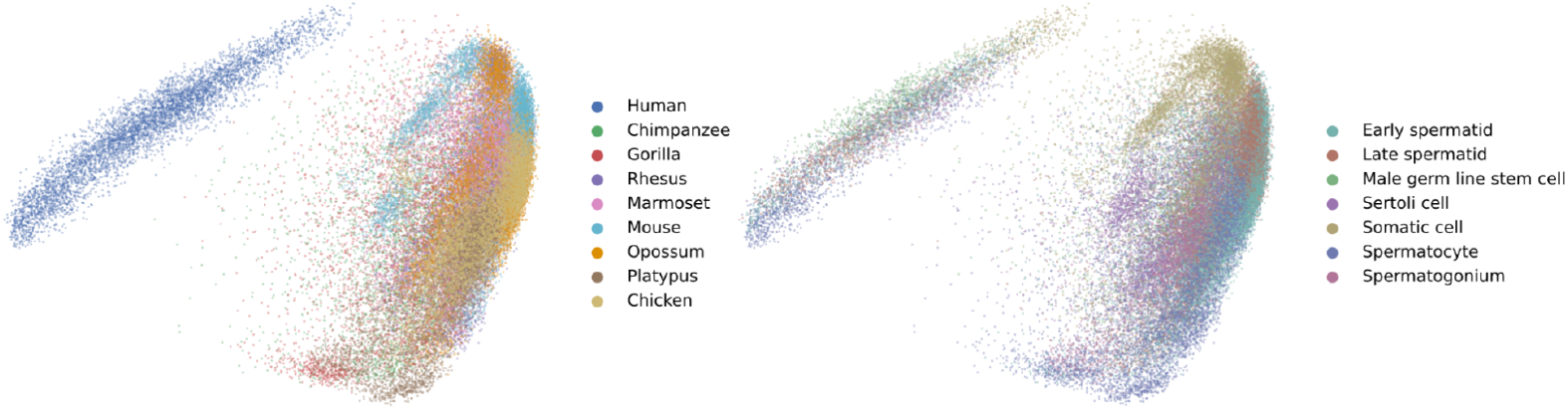
TF-Sapiens embeddings PCA plots on the Spermatogenesis dataset show a pronounced batch effect for human data compared to other species.

**Fig. S3:**
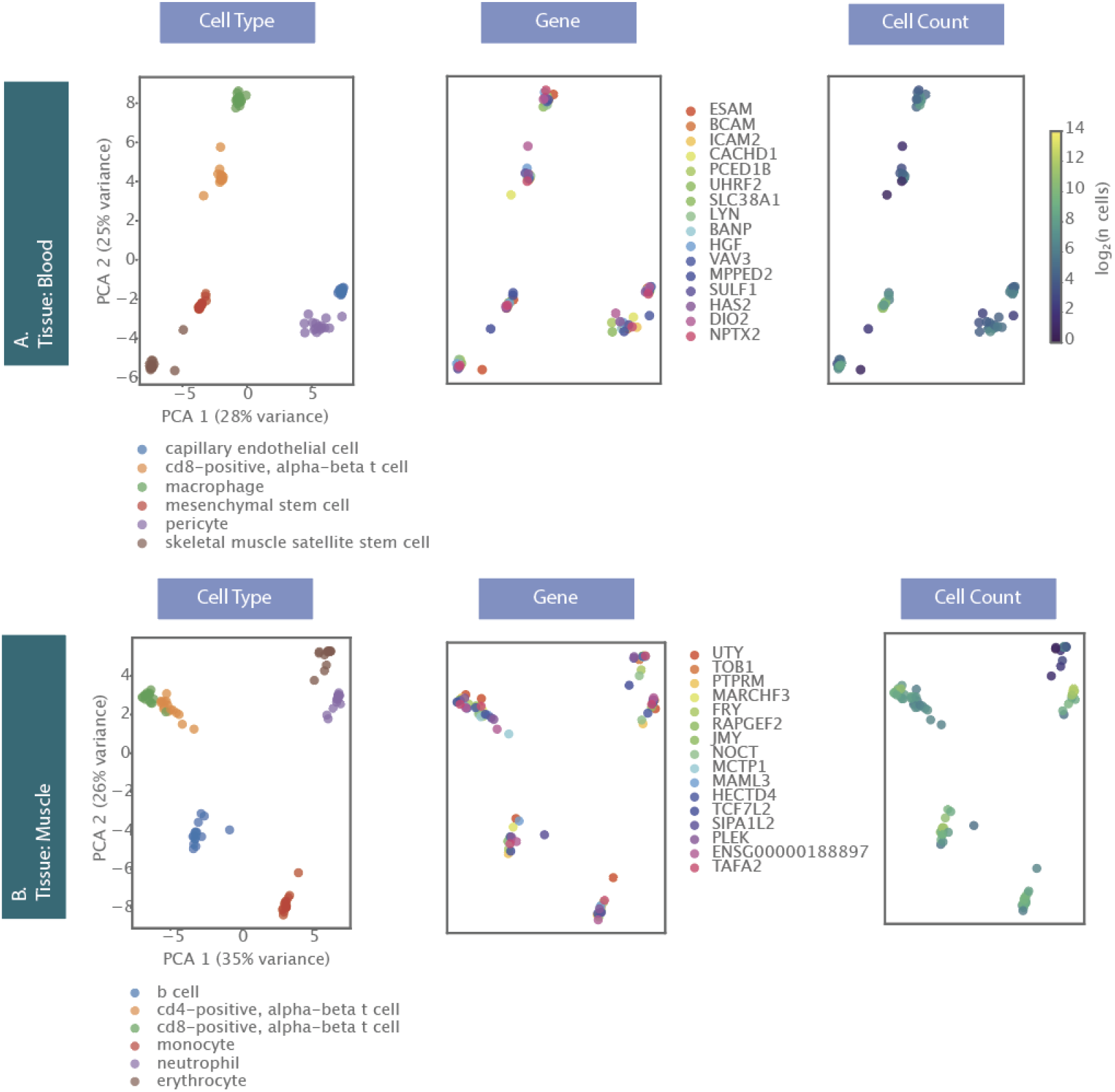
PCA plots of CGE tables constructed per tissue, namely Blood and Muscle from Tabula Sapiens 2. PCA plots are color-coded based on cell type (left column), genes (middle column), and number of cells (log2, right column) used to obtain the average CGE for a given gene. Genes were selected from those highly variable genes in each tissue that are expressed in at least half of cell types and within each cell type are expressed in at least 10% of cells.

**Fig. S4:**
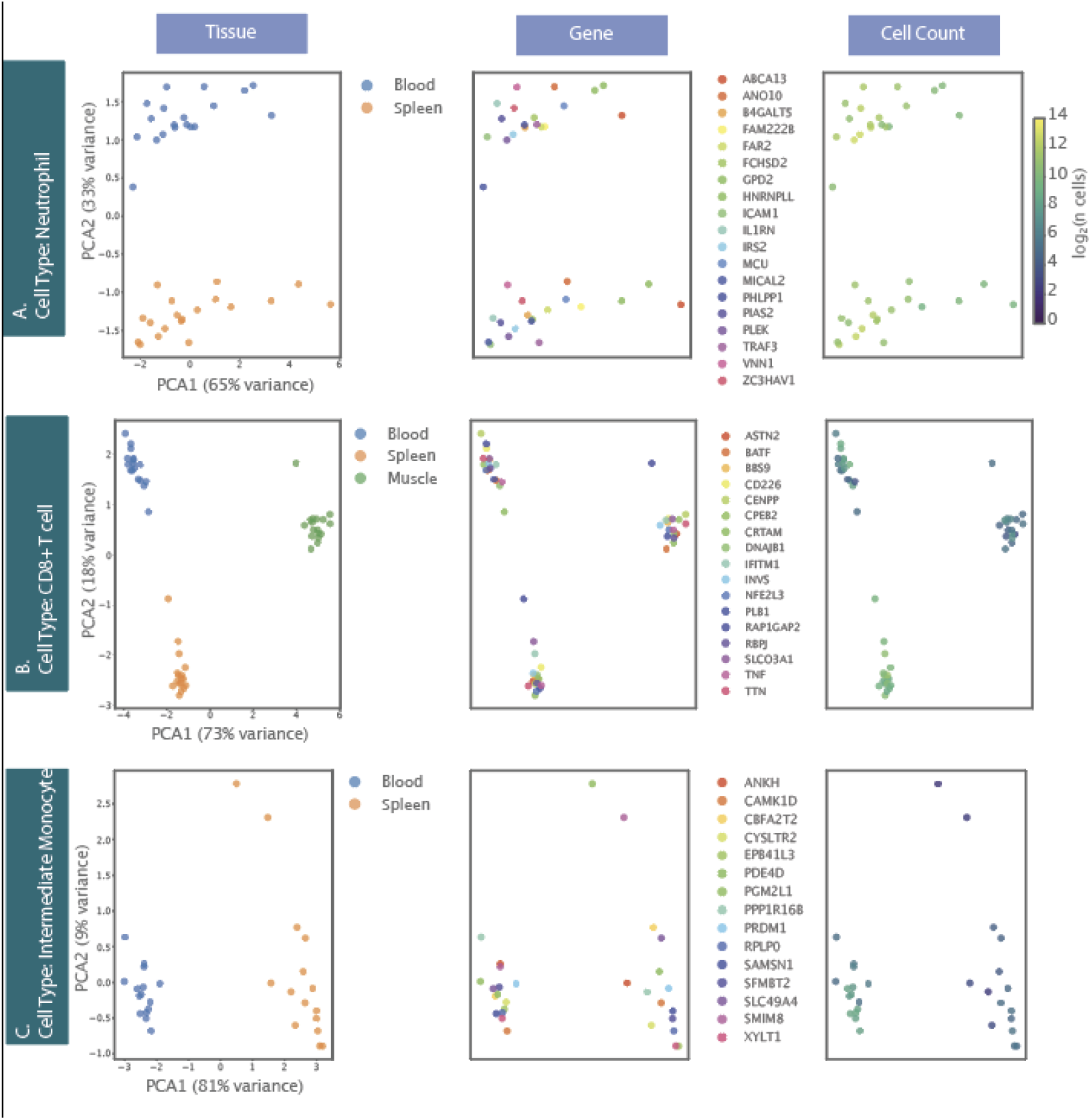
PCA plots of CGE tables constructed per cell type from Tabula Sapiens 2. PCA plots are color-coded based on tissue (left column), genes (middle column), and number of cells (log2, right column) used to obtain the average CGE for a given gene. Genes were selected from those highly variable genes in each tissue that are expressed in at least 10% of cells.

**Fig. S5:**
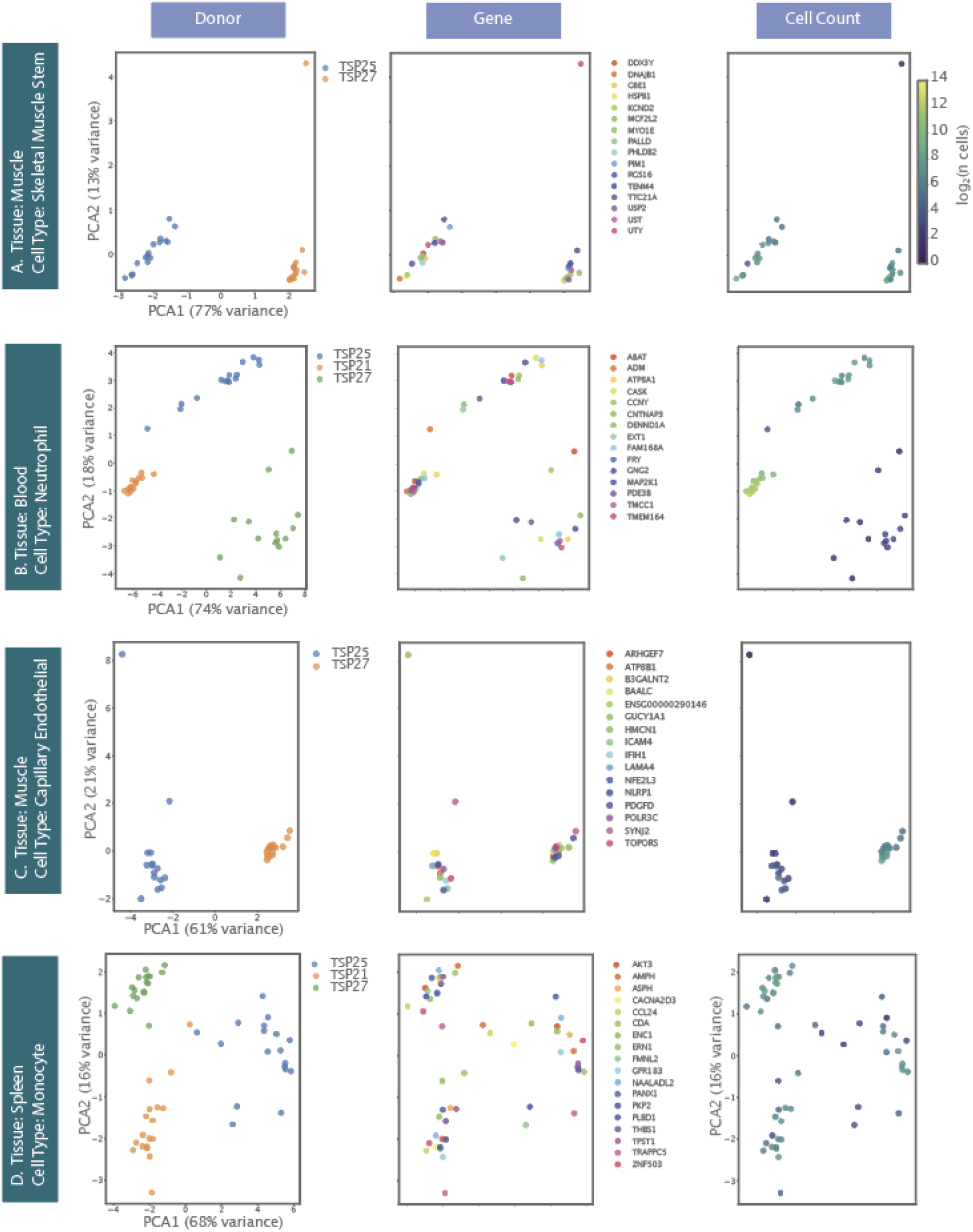
PCA plots of CGE tables constructed per tissue and cell type from Tabula Sapiens 2. PCA plots are color-coded based on donor (left column), genes (middle column), and number of cells (log2, right column) used to obtain the average CGE for a given gene. Genes were selected from those highly variable genes in each tissue cell-type sample, that are expressed in at least at least 10% of cells.

**Fig. S6:**
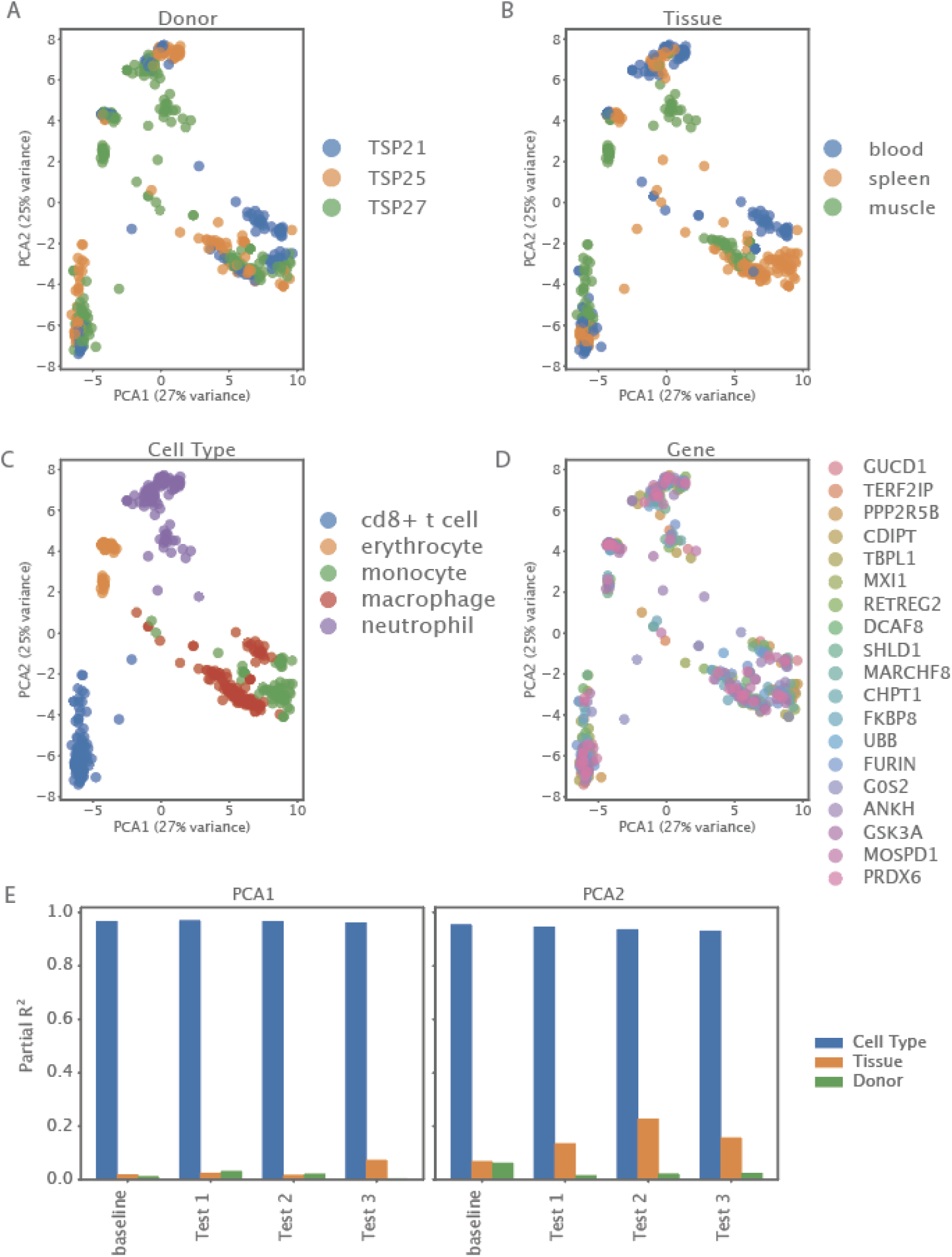
PCA of averaged CGEs color-coded based on donor (A), tissue (B), cell type (C) and gene (D). These results represent the “baseline” gene selection criteria. (E) variance partitioning applied onto the baseline gene filtering criteria as well as three other test cases (see Materials and Methods: CGE).

**Fig. S7:**
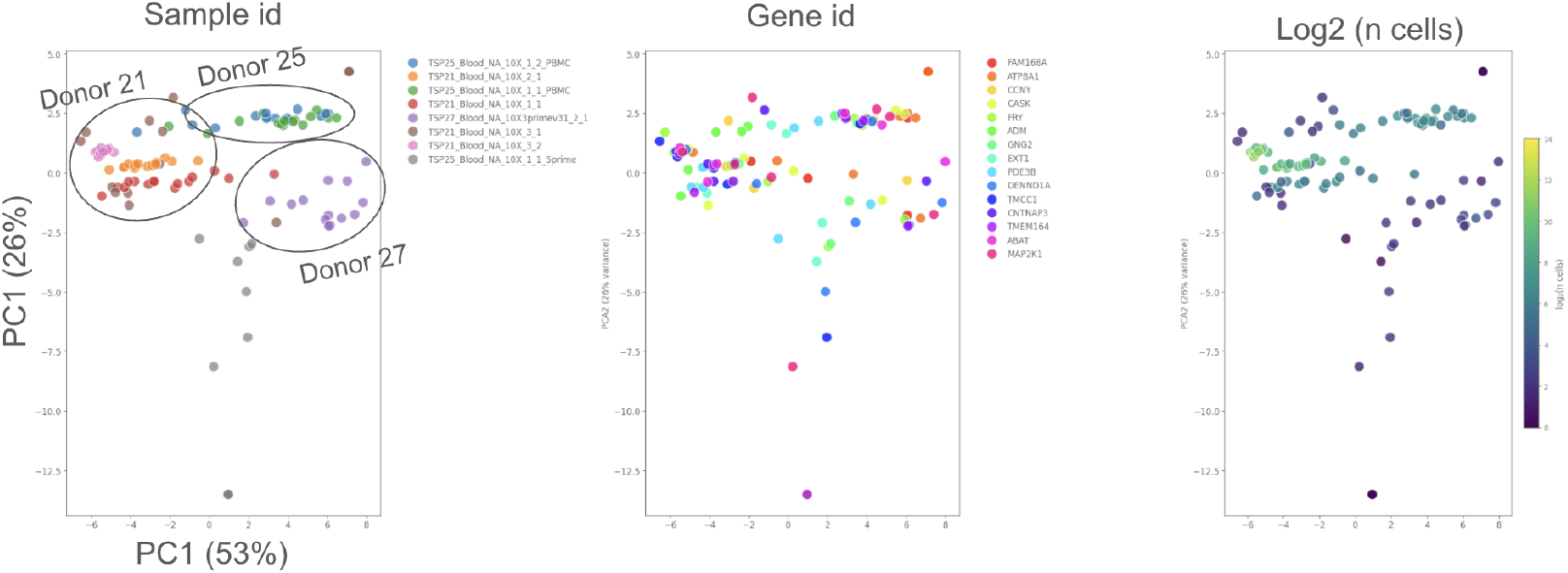
PCA plots of CGE tables constructed per batch from Tabula Sapiens 2 blood neutrophils. PCA plots are color-coded based on batch (left column), genes (middle column), and number of cells (log2, right column) used to obtain the average CGE for a given gene.

**Fig. S8:**
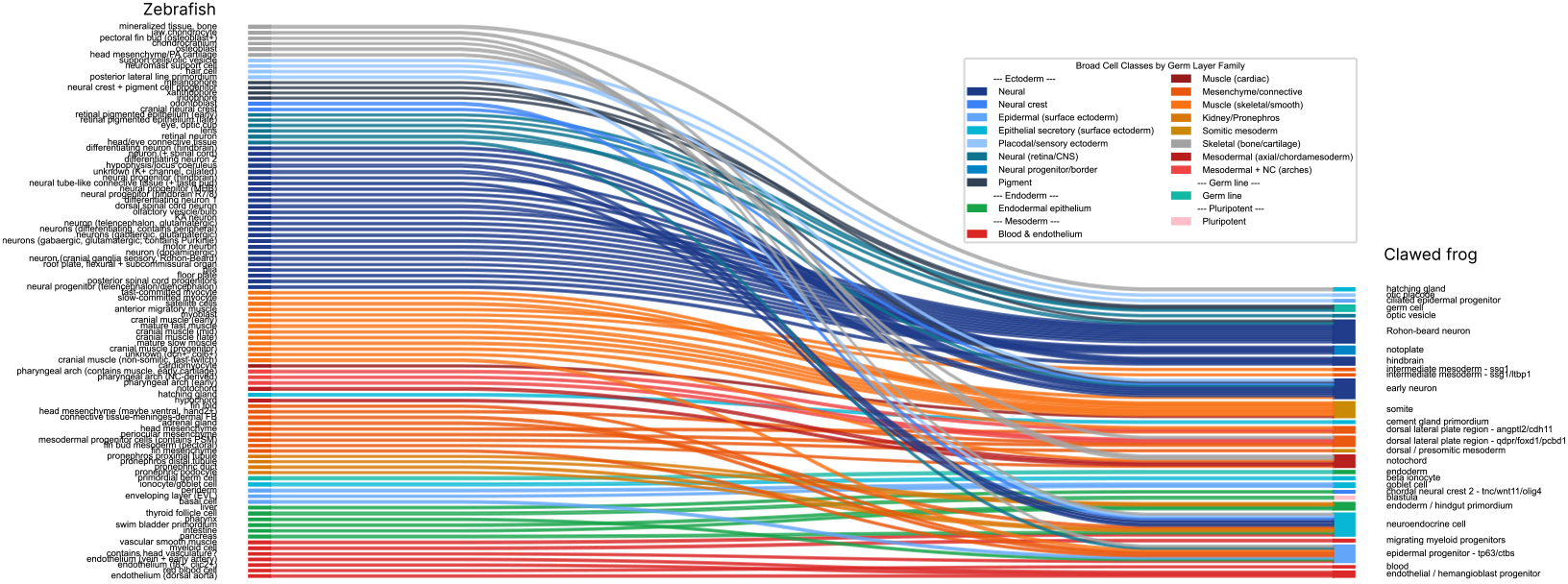
Sankey diagram mapping developmental cell atlases across vertebrate species (zebrafish-clawed frog), colored by cell class and grouped by germ layer.

**Fig. S9:**
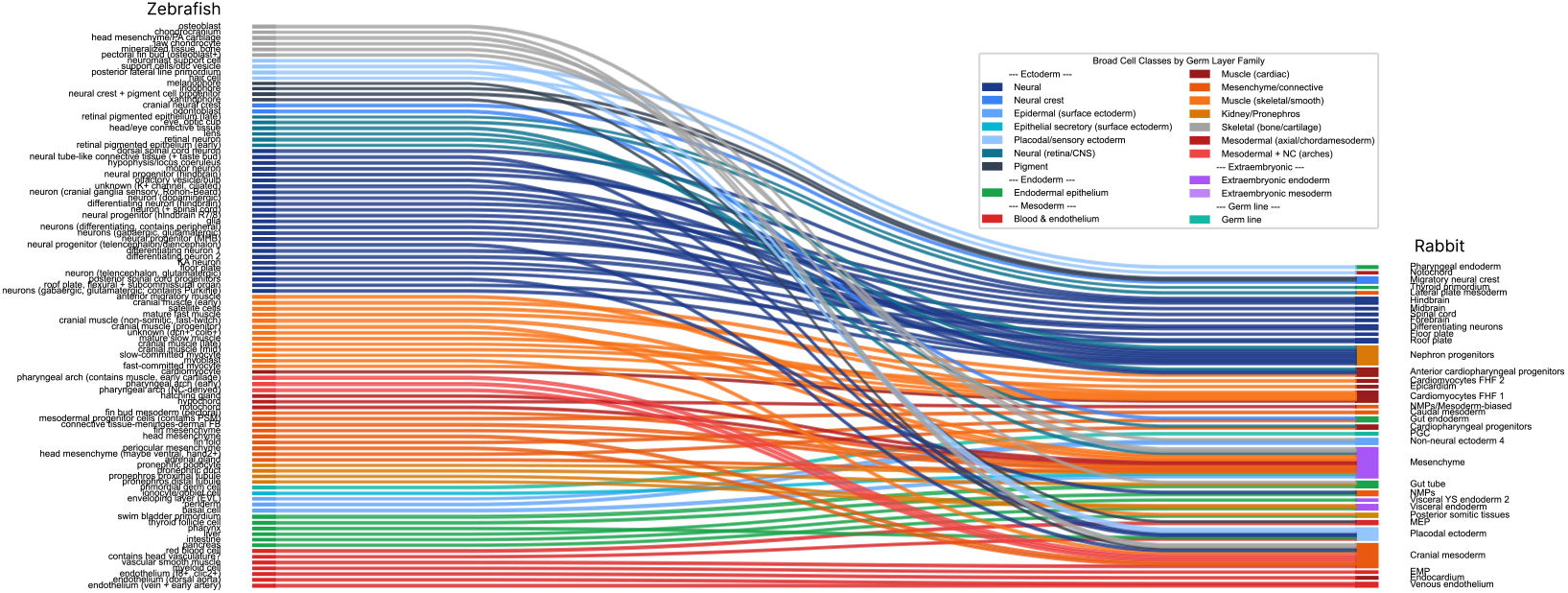
Sankey diagram mapping developmental cell atlases across vertebrate species (zebrafish-rabbit), colored by cell class and grouped by germ layer.

**Fig. S10:**
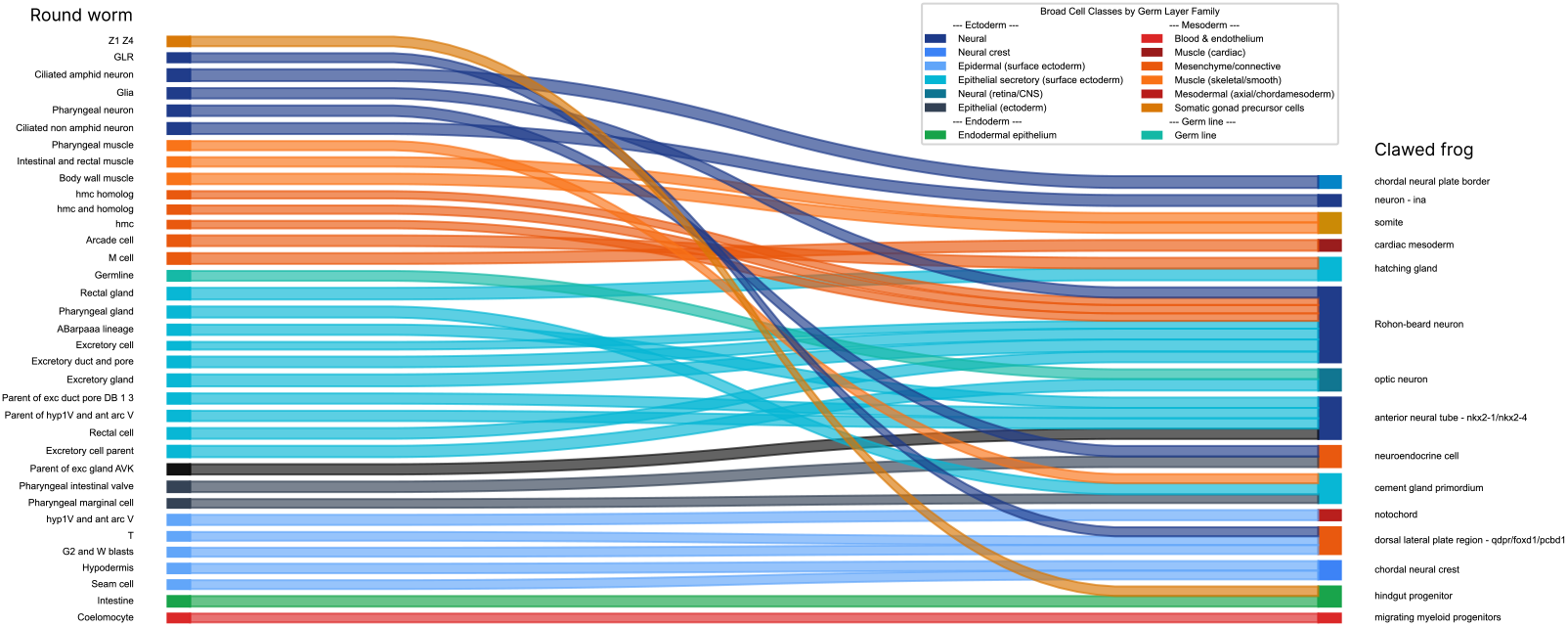
Sankey diagram mapping developmental cell atlases across invertebrate and vertebrate species (*C. elegans*-clawed frog), colored by cell class and grouped by germ layer.

## Notes

### Competing Interest Statement

The authors have declared no competing interest.

### Summary of Updates

Major revisions to the main text including: abstract, introduction, results and conclusion. The revisions follow additional analysis: cross-species drug perturbation, human cell-line drug perturbation and evolutionary cell conservation.

